# AF9/KLF2 gene regulatory circuit links histone lactylation to breast cancer metastasis

**DOI:** 10.1101/2025.06.18.660498

**Authors:** Huida Ma, Ming Yuan, Chenxuan Yang, Yuchen Yuan, Yuanyuan Li, Xin Wang, Zhonghui Tang, Haitao Li

## Abstract

Histone lysine L-lactylation (hereafter referred to as histone Kla) is a novel epigenetic mark induced by glycolytic metabolism, serving as a link between metabolic reprogramming and epigenetic regulation. In this study, we uncover an epigenetic-genetic transcriptional regulatory circuit involving AF9 and KLF2 that drives luminal breast cancer progression. AF9, identified as a reader of histone H3 lysine 9 lactylation (H3K9la), promotes KLF2 expression, while KLF2, functioning as a transcription factor for AF9, forms a positive feedback loop that amplifies lactylation-dependent effects. This circuit activates tumor-associated pathways, including TGFβ1, glucose and lactate transporters, and metabolic enzymes essential for glycolysis and serine biosynthesis, driving tumorigenesis and metastasis. Spatial and single-cell transcriptomics show AF9-positive tumor cells enriched in regions of active lactylation, correlated with immune evasion through interactions with M2 macrophages. Together, AF9, H3K9la, and KLF2 integrate metabolism, epigenetics, and signaling to promote tumor growth and metastasis, highlighting AF9’s central role as a histone lactylation reader and a potential therapeutic target in breast cancer.

**Graphical abstract:** 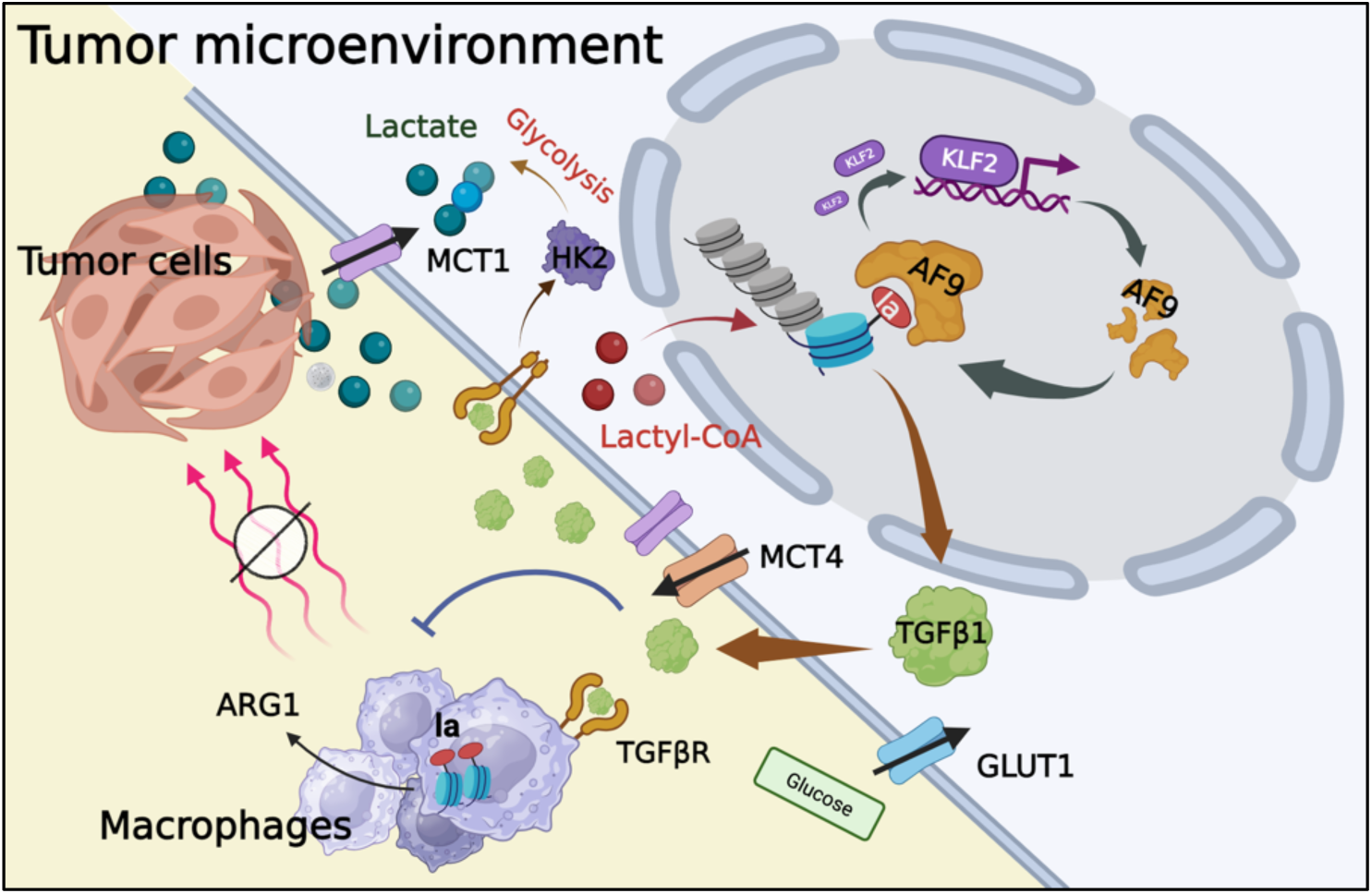

## INTRODUCTION

Histone post-translational modifications (PTMs) are crucial for regulating gene expression and cellular function, linking metabolism and epigenetics and playing significant roles in health and disease^1,2^. L-lactate, a byproduct of glycolysis^3^, is highly upregulated in the tumor microenvironment (TME) under the Warburg effect and hypoxia (L-lactate will be referred to as lactate hereafter for clarity unless otherwise stated)^4,5^. As a major carbon source for cancers, lactate is transported between cells in the TME, and its connection to epigenetics is exemplified by histone lactylation, which plays a critical role in tumor development ^6–8^. For instance, H3K18la has been found to promote the expression of the m6A reader protein YTHDF2, driving oncogenesis in ocular melanoma^9^. Furthermore, the presence of H3K18la has the ability to enhance the expression of METTL3, an enzyme that plays a role in m6A modification^10^. This, in turn, promotes the immunosuppressive capabilities of tumor-infiltrating myeloid cells (TIMs) that are linked to the unfavorable prognosis of colon cancer patients^10^. These findings suggest histone lactylation mediates intercellular communication within the TME.

The regulation of histone lactylation involves both enzymatic mechanisms (CBP/p300 as writers and HDAC1-3 as erasers) and emerging non-enzymatic pathways, which add complexity to its dynamics. ^11–13^ However, the downstream functional response of histone lactylation remains incompletely understood, and the identification and characterization of histone lactylation readers have been a major focus of recent studies. Using combined biochemical, structural, and cell-based approaches, we identified AF9, a YEATS family protein, as a reader of H3K9la, a mark associated with active gene expression. This was likewise confirmed using photoaffinity probe by Li lab^14^. Notably, H3K9la, an active chromatin mark^15^, reduces AF9 binding affinity compared to acetylation, unlike histone crotonylation, which enhances binding^16,17^. However, this reduced binding is compensated by tumor-driven upregulation of AF9 expression via a feedback loop involving the transcriptional factor KLF2. AF9 binds H3K9la at KLF2 promoter regions to drive KLF2 expression, which in turn binds to AF9 promoter to activate its transcription, thus establishing a self-reinforcing circuit. This loop amplifies lactylation-dependent gene expression and drives tumorigenesis pathways, including TGFβ signaling, glycolysis, and serine biosynthesis, ultimately promoting the progression and metastasis of luminal breast cancer.

Through transcriptomic and spatial single-cell analyses of luminal breast cancer tissues, we found significantly elevated levels of AF9, KLF2, and lactylation pathways compared to paracancerous tissues. Given the established role of histone lactylation in regulating anti-inflammatory, immune escape^18^, and wound-healing pathways in macrophages, its upregulation in luminal breast cancer likely contributes to shaping an immunosuppressive TME^11,19,20^. Taken together, our findings reveal AF9, H3K9la, and KLF2 as central players in a metabolic-epigenetic feedback axis that integrates metabolism, epigenetics, and tumor-associated signaling pathways. This feedback axis not only drives tumor growth and metastasis but also highlights the strong dependence of luminal breast cancer on metabolic and environmental signals, providing insight into its high prevalence. Overall, our study establishes luminal breast cancer as a metabolism-associated disease and identifies AF9 and histone lactylation as promising therapeutic targets for disrupting this pathogenic feedback system.

## RESULTS

### AF9 YEATS domain is a histone lactylation reader

To identify readers of histone lactylation (Figure 1A), we purified twelve acylation reader domains, including all human YEATS family proteins (AF9, ENL, GAS41, YEATS2), DPF family proteins (MOZ, MOF, DPF1/2/3, PHF10), and two representative bromodomain (BRD) members (BRD3/9). Binding affinities to corresponding histone peptides were measured using isothermal titration calorimetry (ITC) (Figure S1B). Histone peptides with various acylation sites were chosen based on each reader’s reported sequence preference (Figure S1A)^21^. In eleven cases, histone lactylation disrupted binding to background levels similar to the unmodified form, indicating lactylation is an unfavorable mark for these readers. The exception was the AF9 YEATS domain (Figure 1B), which retained detectable binding to lactylated histone but with compromised affinity compare to acetylated histones. ITC analysis revealed a dissociation constant (*K*_D_) of 74.4 μM for AF9 YEATS binding to H3K9la, approximately 16-fold weaker than its *K*_D_ for H3K9ac (4.6 μM). For site selectivity, the *K*_D_ for H3K18la further dropped to 171.8 μM, while no bindings were detected for H3K14la and H3K27la peptides (Figure 1C). These findings demonstrate that AF9 YEATS domain recognizes histone H3 lactylation, with a site preference for K9 and K18.

**Figure 1.**
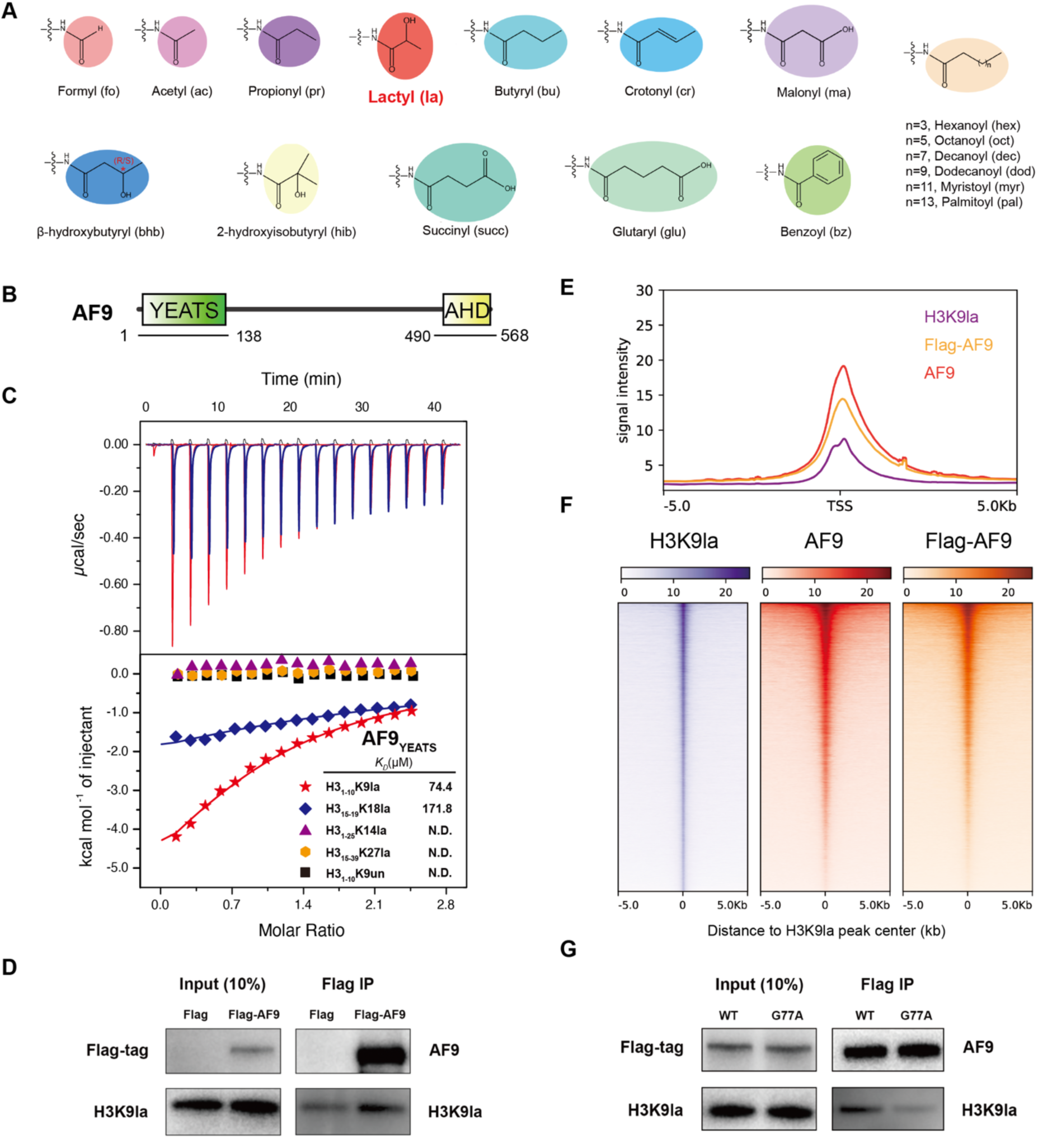
Identification of AF9-YEATS domain as a novel histone lactylation reader. (A) Chemical structures of major lysine acylations. The acylation group is color shaded and lysine lacytlation is highlighted in red. (B) Domain architecture of human AF9. Underlined regions are the YEATS domain of AF9 for binding study^16^ and the AHD domain^42^. Full-length is used to transfection. (C) ITC fitting curves comparing histone lactylation peptides binding preference at sites H3K9, H3K14, H3K18 and H3K27. N.D.: not detectable. (D) AF9 co-immunoprecipitated nucleosomes marked by H3K9la more than control. (E) Average genome-wide occupancies of AF9 (red), Flag-AF9 (orange) and H3K9la (violet) ±5 kb around the TSS in MCF7 cells. (F) Heat maps of the normalized density of AF9 (red), Flag-AF9 (orange) and H3K9la (violet). CUT&Tag tags in MCF7 cells centered on H3K9la-binding peaks in a ±5 kb window. The color key represents the signal density. (G) AF9 co-immunoprecipitated nucleosomes marked by H3K9la in a YEATS-dependent manner. WT indicates the wild type AF9 and G77A is a mutant in AF9 YEATS domain that interrupts the recognition to H3K9la detected in vitro.

To investigate whether AF9 YEATS interacts with H3K9la-modified histones in a cellular context, we conducted co-immunoprecipitation (co-IP) experiments by transfecting Flag-AF9 into MCF7 cells, with an empty vector serving as the negative control (Figure 1D). Immunoprecipitated Flag-AF9 showed a stronger H3K9la signal, indicating its interaction with lactylated histones. Additionally, we analyzed a G77A point mutant that disrupts AF9’s binding to Kla (Figure S2B). Consistent with biochemical findings, H3K9la-modified histones co-immunoprecipitated significantly more with wild-type AF9 than with the G77A mutant (Figure 1G). These findings confirm that AF9 recognizes H3K9la in vivo and that this interaction is mediated by the AF9 YEATS domain.

To examine AF9 chromatin occupancy and its correlation with H3K9la, we performed CUT&Tag experiments followed by high-throughput sequencing. Using anti-Flag M2 and anti-AF9 antibodies, we identified 38,583 and 38,253 AF9-enriched peaks, respectively, with strong enrichment in promoter regions (±5 kb around the transcription start site, TSS) (Figure S6A, S6B and 1E). H3K9la CUT&Tag revealed 44,249 peaks in MCF7 cells, with over 50% of AF9 and Flag-AF9-occupied genes overlapping H3K9la-enriched regions. Heat map analysis further confirmed the co-localization of AF9 with H3K9la (Figure 1F). These results show that AF9 recognizes H3K9la at promoter regions to facilitate gene activation.

### Molecular basis underlying AF9 in recognition of histone lactylation

To gain molecular insights into the reader function of the AF9 YEATS domain, we solved the crystal structure of human AF9 YEATS domain (1-138) bound to the H3_1-10_K9la peptide at 2.9 Å resolution. The overall engagement mode of H3K9la peptide with AF9 YEATS domain was similar to H3K9ac peptide. AF9 YEATS adopts an immunoglobin (Ig) fold, consisting of a two-layer β-sandwich formed by eight antiparallel β-strands (Figure 2A). Based on the electron density map, we could clearly build the H3_4-10_K9la peptide. The H3 binding surface is negatively charged, thereby electrostatically facilitating the recognition of the basic H3 peptide (Figure 2B). The side chain of H3K9la is positioned in the same pocket as H3K9ac readout, with the lacylation side chain sandwiched by two aromatic residues F59 and Y78 (Figure S2A). Due to the branched non-flat feature of lactyl group, the binding is not as snug as acetylation. In comparison to acetylation, lactylation results in an expansion of the binding pocket (Figure 2D). This expansion causes varying degrees of outward displacement in the residues S58, F59, and Y78, which could explain the decreased binding affinity observed in lactylation (Figure S1B). In the complex structure, AF9 YEATS forms an extensive hydrogen bonding network with the H3K9la peptide, involving direct (Figure 2C) or water-mediated interactions conducted by the key residues shown in Figure S2A. Mutagenesis studies confirmed that Alanine substitution of H56, F59, G77 and D103 completely abolished AF9 YEATS and H3K9la binding (Figure S2B). Interestingly, we found that S58R, but not S58A, totally lost its H3K9la binding ability, whereas S58T enhanced the interaction (Figure S2C), indicating that the small and polar feature of S58 is crucial for H3K9la recognition (Figure 2C).

**Figure 2.**
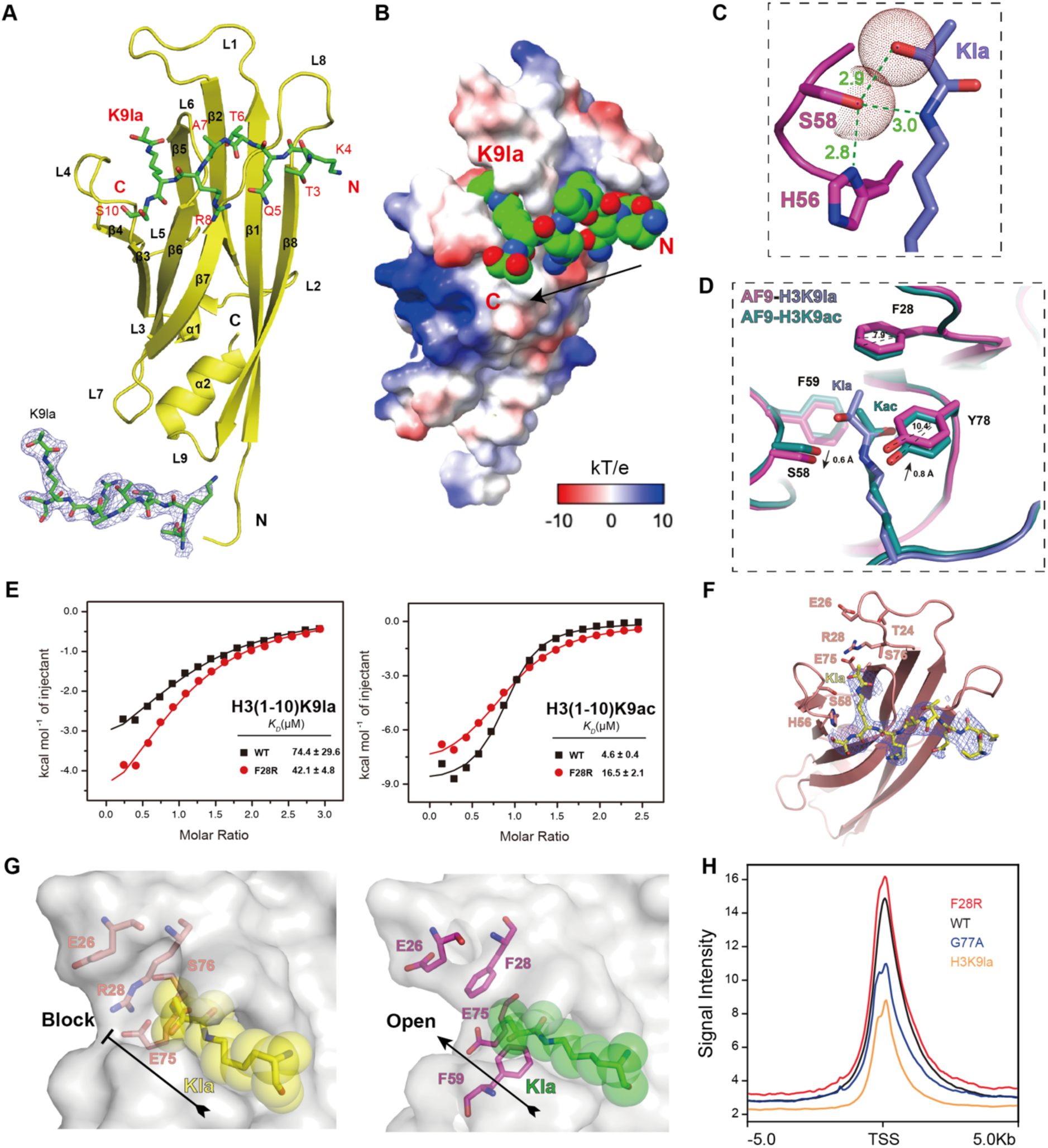
Structural basis for histone Kla recognition by AF9 YEATS domain. (A) Overall structure of AF9 YEATS domain bound to H3K9la peptide (PDB ID: 8Z73). AF9 YEATS is shown as yellow ribbon. H3K9la peptide is shown as green sticks covered by simulated annealing 2Fo-Fc omit map contoured at 1.0σ level. (B) The electrostatic surface view of the AF9YEATS-H3K9la complex structure. Electrostatic potential is expressed as a spectrum ranging from -10kT/e (red) to +10kT/e (blue). The H3K9la peptide is depicted as space-filling sphere with green for carbon, blue for nitrogen and red for oxygen atoms. (C) Stereo view of hydrogen bonding network involving H3 side chain (green sticks) and residues in AF9 YEATS (yellow sticks). Red dotted lines, hydrogen bonds; Cyan balls, water molecules. (E) ITC assays of H3K9la (left) and H3K9ac (right) binding by WT or F28R mutant of AF9 YEATS domain. (F) Overall structure of AF9-F28R YEATS domain bound to H3K9la peptide (PDB ID: 9IM4). H3K9la peptide is shown as green sticks covered by simulated annealing 2Fo-Fc omit map countered at 1.0σ level. (G) Close-up view of the K9la-binding pocket of the WT (right) and F28R (left) AF9 YEATS domain. The pocket is displayed as semi-transparent surface with key residues shown as sticks. Kla is depicted in both stick and space-filling sphere modes. (H) Metaplots depicting the co-localization of Flag-AF9 WT (black), F28R(red), G77A (blue) and H3K9la (orange) in gene promoters.

The other human YEATS proteins (including ENL, GAS41 and YEATS2) showed no detectable histone lactylation recognition in our ITC assay. Their binding towards histone acetylation are much weaker than that of AF9 YEATS (Figure S1B), which might explain their even weaker so as to undetectable association with Kla. Based on both sequence^22^ (Figure S2G) and structure (Figure S2H and S2I) analysis, we found that in our reported AF9 structure^23^, H3K4 formed a stacking with the imidazole rings of AF9 H107 and H111 residues, thereby strengthening their interactions. Notably, these two positions in ENL are Asn instead of His, indicating a lacking of the stackings in this region of ENL (Figure S2H). To verify this, the AF9 YEATS mutant H107H111 to N107N111 (indicated as HHNN) reduced the H3K9la-binding affinity by 1.5fold (Figure S2B). As for GAS41 and YEATS2 YEATS domains, we observed that they have a Tyr residue at position 59, which forms hydrogen bonds with a water molecule and has a greater tendency to recognize planar modifications rather than branched modifications (Figure S2I). In the structure, mutating F59 to Y indeed results in a steric clash with the lactyl group (Figure S2F). To confirm this, we substituted the F59 with Tyr in AF9 (F59Y) and performed ITC titration experiment. The results showed that this F59Y mutation diminished AF9’s affinity for H3K9la by 2-folds (Figure S2B). Notably, both GAS41 and YEATS2 YEATS domains utilize an antiparallel histone binding mode to that of AF9 YEATS, which may account for their relatively low affinities (Figure S2I). These features collectively make AF9 unique to be able to recognize histone lactylation among human YEATS proteins.

### Key Contribution of H3K9la in AF9-mediated Transcriptional Regulation

The specific role of H3K9la in AF9 function was further explored through the design of an AF9 F28R mutant and CUT&Tag experiments. From the CUT&Tag experiment, we observed the co-localization of H3K9la and H3K9ac (Figure S4A and S4B). In vivo, the abundance of H3K9ac is higher than that of H3K9la (Figure S3). For further functional study, we designed an AF9 mutant (F28R) that could enhance the recognition of H3K9la but reduce H3K9ac binding (Figure 2E). Co-IP experiments further verified the functional specificity of this mutant, supporting the unique role of H3K9la in transcriptional regulation.

To explore the underlying mechanism, we then solved the AF9-F28R YEATS domain structure in complex with H3K9la at 2.8 Å resolution (Figure 2F). The reinforcement could attribute to the electrostatic interaction between F28R and E75, forming a polar gate to accommodate the side chain (Figure S2E). As a result of this, F28R mutant formed “electrostatic encapsulation” at the end of the binding pocket, impacting the stretch of lactyl-lysine side chain (Figures 2G). By CUT&Tag experiments, we further validated that the F28R mutant of AF9 exhibits higher enrichment at the TSS region compared to the WT, significantly higher than the G77A binding-disruption mutant (Figure 2H).

Although H3K9la and H3K9ac co-localize on the genome, our results revealed that H3K9la plays a critical, non-redundant role in guiding AF9 to target gene promoters. These observations highlight the idea that histone modifications within the same chromatin region exist exclusively at a given time, and when H3K9la is present, even transiently, it is indispensable for regulatory events. H3K9la’s event occupancy during these transitions is effectively 100%, signifying its pivotal role as an “on-demand” epigenetic signal. This logic parallels ligand-induced signaling in pathways such as those mediated by the epidermal growth factor receptor (EGFR). EGFR is phosphorylated on its intracellular domain only upon ligand binding, enabling downstream signaling^24–26^. Once the ligand dissociates, phosphatases rapidly remove the phosphorylation. Despite the short-lived nature of the modification, its presence during the signaling event is absolute and critical. Similarly, H3K9la induces regulatory changes by guiding AF9 to transcriptionally active promoters, contributing to gene expression switches in a manner that cannot be captured by its co-occurrence with H3K9ac. Thus, our findings demonstrate that H3K9la is not merely a passive modification but actively orchestrates AF9 localization and transcriptional control, serving as a key signal for epigenetic regulation.

### AF9-H3K9la recognition promotes luminal breast cancer development

The mutation and amplification of AF9 is closely linked to various human cancers, especially its abnormal amplification in breast cancer (Figure 3A). Based on the analysis of TCGA data sets, we found that AF9 was significantly increased in luminal A breast cancer patients and associated with poor prognosis (Figures S5A and S5B). We thus deduced that AF9 was linked to the development of luminal breast cancer tumors. This was confirmed through western blot detection in both luminal A and B breast cancer patients. AF9 expression is much higher in tumor tissue than in adjacent normal tissue (Figure 3D).

**Figure 3.**
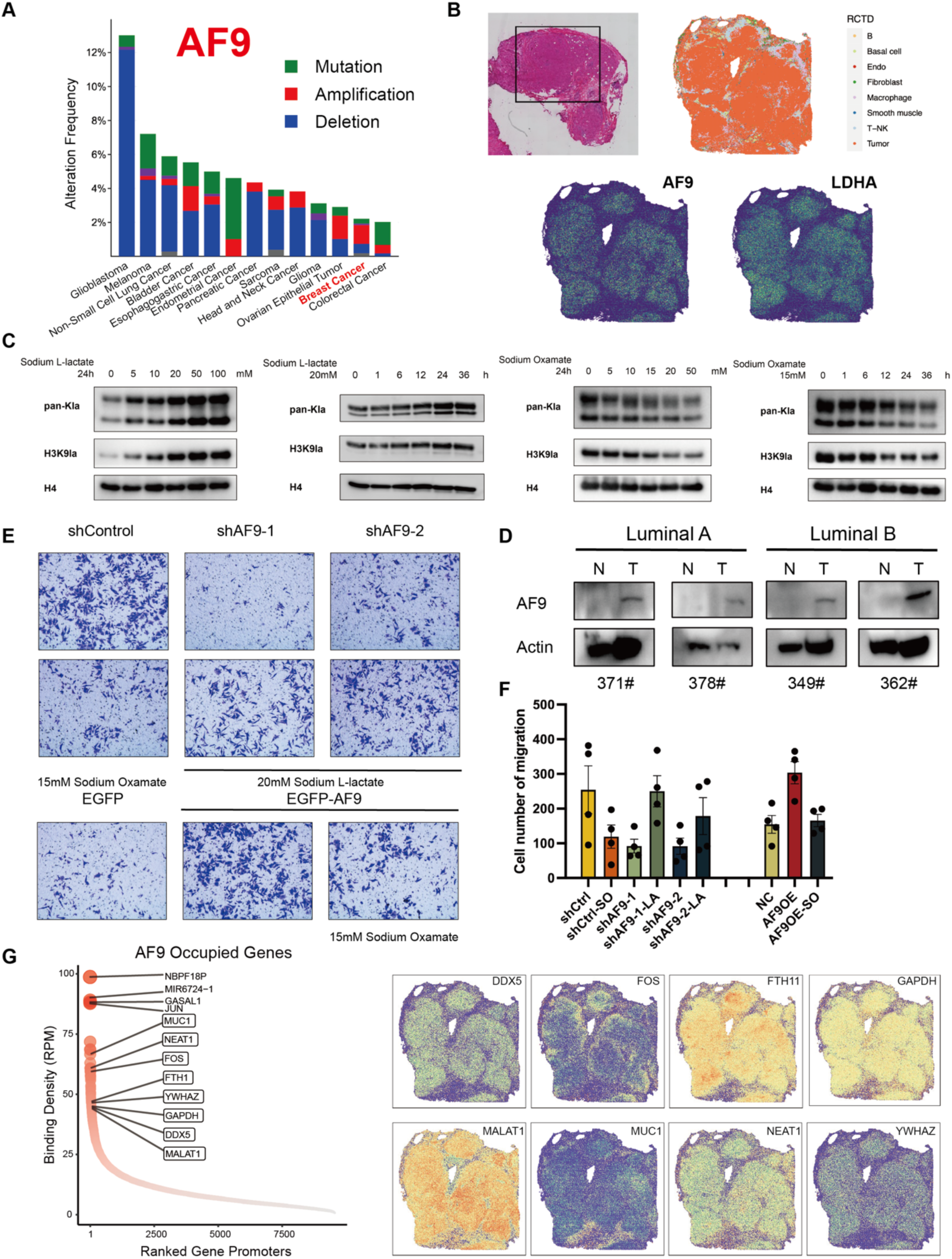
Histone lactylation is an inducible epigenetic mark and required for breast cancer development. (A) Histogram showing the alteration frequency of AF9 in human cancers. Data was acquired from cBioPortal. (B) Visualization of the distribution of AF9 and LDHA expression across tissue sections from luminal breast cancer patient. The picture above shows HE staining and cell type distribution. (C) Western blot analysis of histone Kla and H3K9la levels in response to the indicated time and concentration of sodium L-lactate and sodium oxamate gradient treatment in MCF7 cells. (D) AF9 is amplified in the tumor compared with paratumor samples in luminal breast cancer patients. (E) AF9 is required for MCF7 cell invasion. Transwell assays of cells were stained with crystal violet 42h after seeding. (F) The results of transwell are quantified and statistically analyzed through invasion experiments by ImageJ. (G) The map shows AF9 localized genes arranged in order of intensity (left). Many of the AF9 strongly localized genes are highly expressed in tumor cells (right).

To better understand the distribution of AF9 expression in luminal breast cancer, we used spatial transcriptomic technology and found that AF9 is highly expressed in tumor cells (Figure 3B). LDHA, the lactate dehydrogenase, shows a similar pattern to AF9, indicating a potential correlation between lactate and tumor cells (Figure 3B). We then performed immunoblotting using pan-Kla and AF9 antibody in multiple mammalian cell lines with or without sodium L-lactate treatment and analyzed the levels of histone lactylation and AF9 (Figure S5D and S5E). The high concentration of sodium L-lactate treatment significantly increased the levels of AF9 and pan-Kla in MCF7, a common human luminal breast cancer cell line (Figure S5D).

In order to further verify the impact of AF9-Kla on tumor characteristics at the cellular level, we constructed the stable transfection MCF7 cell line with AF9 knockdown and overexpression (Figure S5C). It has been reported that elevated level of lactate is correlated with metastatic potential in various human primary carcinomas^27^, probably serving as the metabolic precursor of histone lactylation. By treating cells with sodium L-lactate at different concentrations and time points, we demonstrated that histone Kla is inducible (Figure 3C). Our treatment assays identified the optimal treatment duration (24 h) and concentrations of sodium L-lactate (20 mM) and sodium oxamate (15 mM). Subsequently, we examined alterations in tumorigenic properties of MCF7 cells following AF9 knockdown or sodium lactate treatment. In colony forming assays, AF9 knockdown cells formed fewer colonies than control cells, a phenotype that resembled the effect of sodium oxamate treatment (Figure S5F). Similarly, in transwell invasion assays, AF9 knockdown impaired cell invasiveness, mirroring the effect of sodium oxamate; notably, this reduction in invasiveness could be rescued by sodium L-lactate treatment (Figure 3E and 3F). These results indicated that AF9-Kla is required for MCF7 cell proliferation and invasion. Notably, many AF9-localized genes showed elevated expression in malignant cells (Figure 3G), implying that AF9’s ability to recognize histone lactylation marks may serve as an epigenetic mechanism regulating oncogenic gene networks.

### AF9 regulates the metabolism associated genes in MCF7 cells

Given the opposite patterns of MCF7 and H1299 in cell line screening experiments (Figure S5D), we detected the metabolomics of MCF7 and H1299 cell, showing a divergent metabolic state (Figure 4A). MCF7 cells exhibited a greater reliance on glycolytic metabolism and a reduced dependence on TCA cycle metabolism in comparison to H1299 cells. Single-cell sequencing analysis of breast cancer patients identified AF9-positive epithelial cells as displaying enhanced glycolytic activity and nitrogen metabolism, both of which are essential for tumor progression (Figure 4B). The results implied that AF9-Kla recognition axis plays a critical role in mediating metabolic reprogramming in cancer cells.

**Figure 4.**
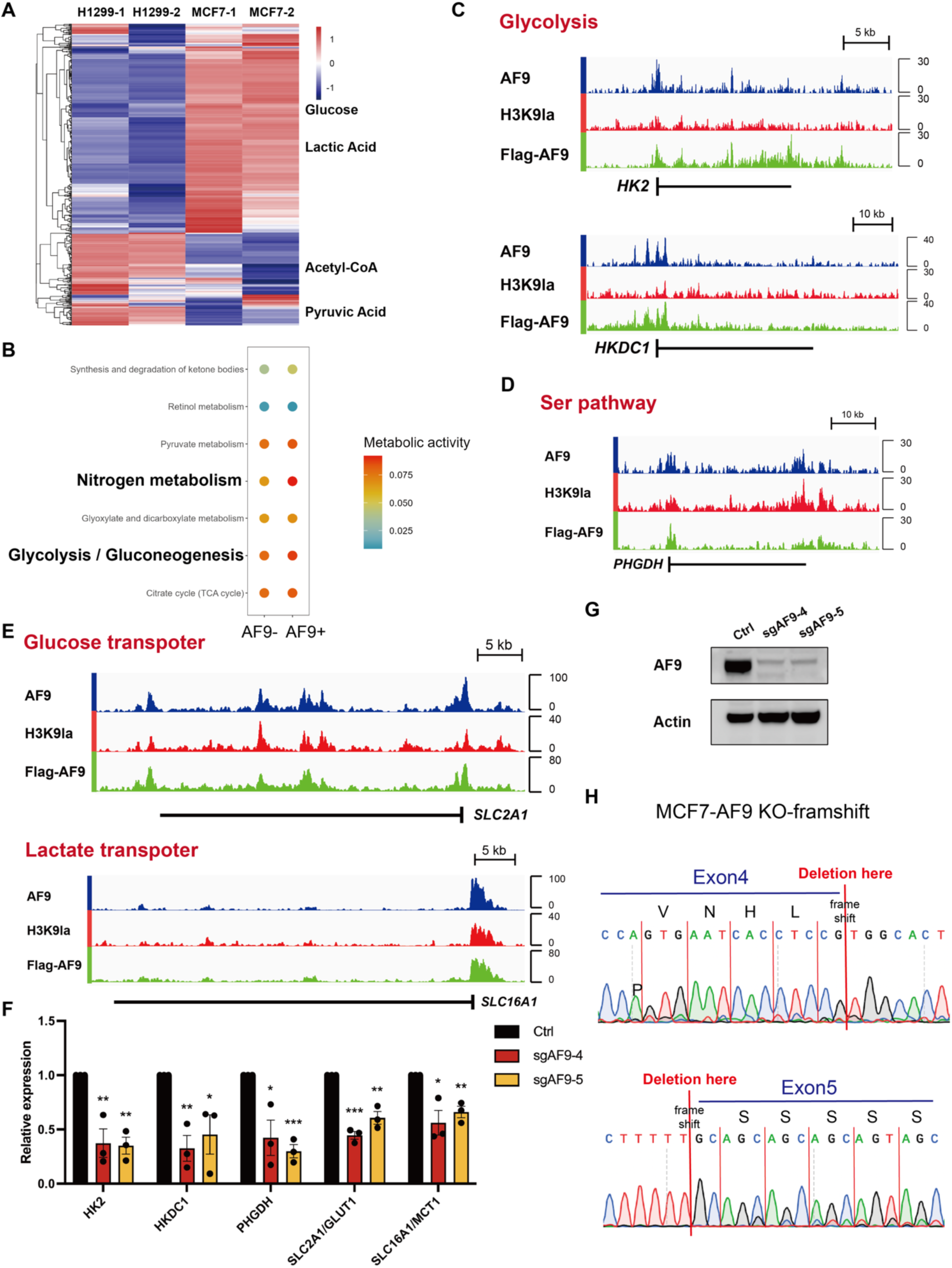
AF9 regulates the metabolic genes expression associated with tumorigenesis. (A) Heat map displaying metabolites detected in H1299 and MCF7 cells. (B) Metabolic pathway analysis of AF9-positive and -negative cells in single-cell sequencing data. (C-E) Genome browser view of AF9 (blue), H3K9la (red) and Flag-AF9 (green) CUT&Tag peaks on the indicated metabolic genes in MCF7 cells. (F) RT-qPCR analysis datas of metabolic genes in AF9-KO MCF7 cells. (G) Western blot to show the knock-out of AF9 in MCF7 cells. (H) Sanger sequencing to show frame-shifting mutation and knock-out of AF9 in MCF7 cells.

Based on the CUT&Tag experiments, we screened potential downstream genes. Genome-wide mapping revealed that both AF9 and H3K9la specifically localize to promoter regions of key glycolytic enzymes, including HK2 and HKDC1 (Figure 4C, S4C and S4D). It was noted that HK2 expression was necessary for the induced gene expression through histone lactylation, but not through histone acetylation^28^. Meantime, we also identified co-occupancy of AF9 and H3K9la at PHGDH (Figure 4D and S4E), which was identified as a significant gene in breast cancer, and reducing its expression resulted in decreased weight of allograft tumors^29^. Furthermore, metabolic transporter genes, SLC2A1 (GLUT1) and SLC16A1 (MCT1), were similarly found to contain AF9 and H3K9la marks in their promoter regions (Figure 4E). Following AF9 knockout in MCF7 cells (Figure 4G and 4H), we quantified expression changes in the metabolically relevant genes identified at AF9/H3K9la-occupied loci. AF9 was found to enhance the expression of glucose transporter GLUT1 (Figure 4F). This suggests that AF9 can identify H3K9la and stimulate glycolysis, leading to increased lactate production. Furthermore, we identified potential regulation of other transporter genes by this recognition pair. Even though all MCTs are bidirectional symports, MCT4 mainly facilitates lactate export while, MCT1 plays a key role in cellular lactate uptake^27^. AF9 was able to upregulate the lactate transporter MCT1, resulting in an increase in lactate content (Figure 4F). The epigenetic-metabolic crosstalk between histone lactylation and lactate transport creates a self-reinforcing cycle that may contribute to immune escape tumor microenvironments through lactate accumulation.

### The AF9-H3K9la recognition pair promotes TGFβ1 expression

To investigate the functional role of AF9-Kla recognition in breast cancer progression, we conducted pathway enrichment analysis comparing AF9-positive and AF9-negative epithelial cells using single-cell RNA sequencing data (Figure 5A). This analysis revealed significant enrichment of TGFβ signaling pathway activity in AF9-positive populations (Figure 5B), suggesting a potential mechanistic link between histone lactylation recognition and TGFβ-mediated oncogenic processes. Consistent with the single-cell results, KEGG pathway enrichment of AF9 CUT&Tag targets similarly identified the TGFβ signaling pathway as enriched (Figure 5C). This suggests that AF9 is correlated to the TGFβ signaling pathway, and we hypothesize that this association is due to the ability of AF9 to regulate gene expression within these pathways.

**Figure 5.**
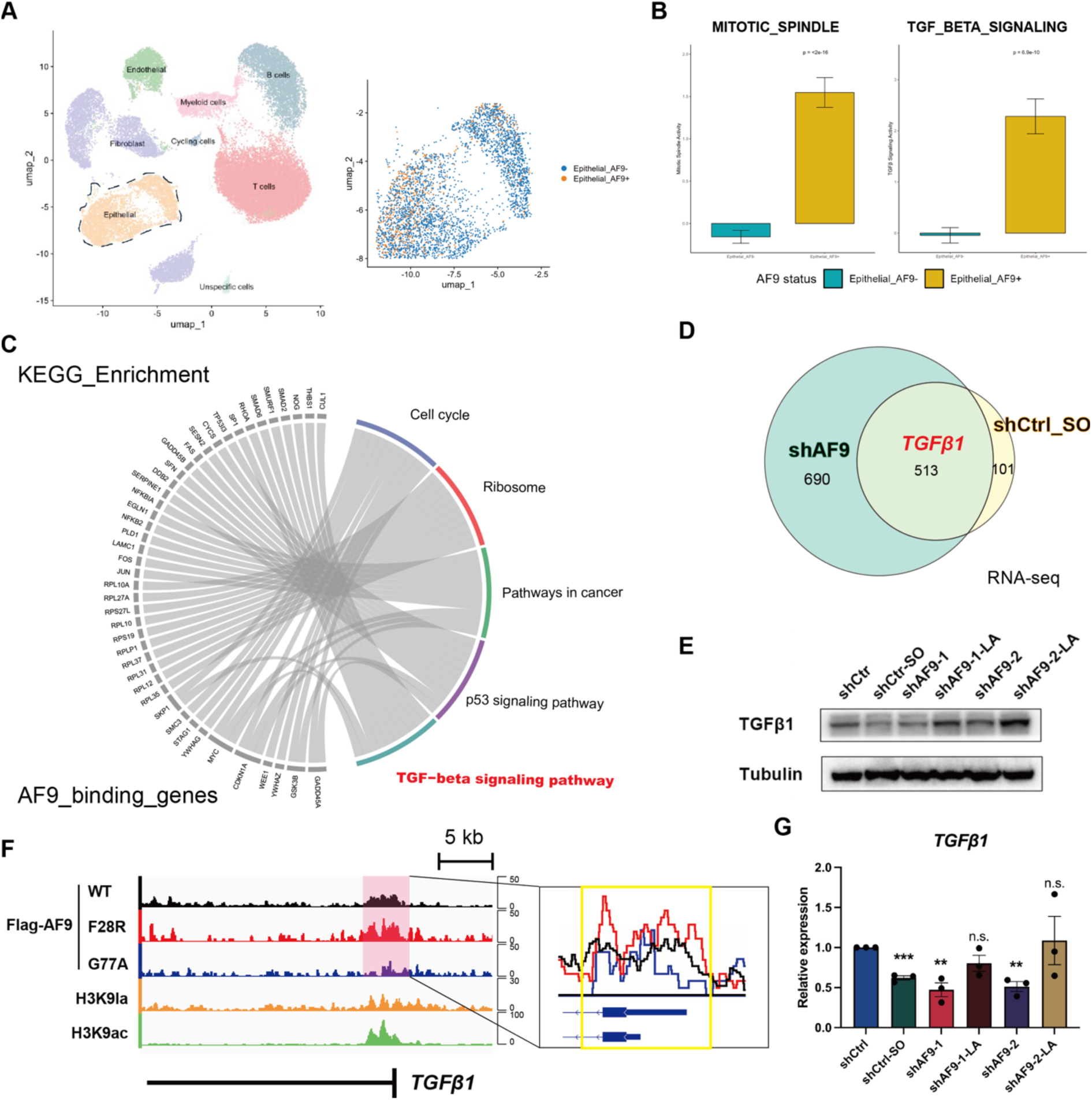
AF9 is recruited to TGFβ1 promoters and correlates with H3K9la. (A) Cell grouping of single-cell sequencing data, dividing epithelial cells into AF9-positive and -negative categories. (B) Pathway analysis finds stronger TGFβ pathway in AF9-positive cells. (C) The KEGG pathways enrichment analysis of AF9 binding genes. (D) The Venn diagram shows the RNA-seq data of AF9 knockdown and sodium oxamate treatment in MCF7 cells. (E) Western blot analysis of TGFβ1 in AF9 knockdown MCF7 cells. (F) Genome browser view of the Flag-AF9WT (black), Flag-AF9F28R (red), Flag-AF9G77A (blue), H3K9la (orange) and H3K9ac (green) peaks on the indicated genes of TGFβ1 in MCF7 cells. Close-up views, positioning the Flag-AF9 WT and mutant peaks located at the promoter regions. (G) RT-qPCR analysis of TGFβ1 in AF9 knockdown MCF7 cells. (∗) P < 0.05; (∗∗) P < 0.01; (∗∗∗) P < 0.001, two-tailed unpaired Student’s test.

To systematically identify genes regulated by AF9 and histone lactylation, we performed RNA-seq analysis comparing AF9-knockdown and sodium oxamate-treated MCF7 cells. Among these, TGFβ1 emerged as a particular candidate given its well-established roles in oncogenesis (Figure 5D). Likewise, statistical analysis of single-cell sequencing data revealed a positive correlation between TGFβ1 and AF9 (Figure S6C and S6D). TGFβ1 is a well-established protein that has been show to promote breast cancer local invasion and liver metastasis^30^. Spatial profiling demonstrated TGFβ1 enrichment in peritumoral regions (Figure S6F). Notably, the expression of TGFβ2, which is another ligand of the TGFβ signaling pathway, could be induced by lactate in glioma cells^31^. Similar to the regulation of TGFβ2, we found that AF9 knockdown reduced TGFβ1 expression, an effect similarly observed with sodium oxamate treatment (Figure 5E and 5G). The observed correlations indicate that AF9 may modulate TGFβ signaling in cancer cells through transcriptional control of TGFβ1.

To further elucidate the regulatory connection between TGFβ1 expression and the AF9-H3K9la reocognition, we conducted CUT&Tag sequencing in MCF7 cells to comprehensively map AF9 chromatin occupancy and its spatial correlation with H3K9la modifications. As expected, TGFβ1 has the localization of AF9 and H3K9la (Figure 5F). When we transfected Flag-tagged AF9-G77A mutant that could interrupt the binding event of AF9 YEAST-H3K9la, the occupancy of AF9 on TGFβ1 declined sharply, suggesting that the AF9-Kla interaction positively regulated gene expression in a YEATS-dependent manner (Figure 5F). Concurrently, we sought to discriminate between AF9-mediated gene activation through Kla recognition versus Kac binding, given the genome-wide cooccurrence of these histone modifications. We transfected cells with Flag-tagged AF9-F28R mutant, which could reduce the recognition of AF9-H3K9ac while boosting the recognition of AF9-H3K9la. After normalization, we found that the signal peak strength of Flag-AF9-F28R was stronger than Flag-AF9-WT on AF9-H3K9la associated genes such as TGFβ1, indicating the main contribution of AF9-H3K9la recognition (Figure 5F). The findings indicated that H3K9la primarily facilitated the recruitment of AF9 to the TGFβ1 gene promoters, subsequently impacting tumor progression via the TGFβ signaling pathway.

### AF9/H3K9la/KLF2 form a positive feedback loop in MCF7

Remarkably, AF9 expression displayed temporal dynamics closely paralleling Kla accumulation during sodium L-lactate treatment. To validate this relationship, we employed dose-dependent treatments with sodium L-lactate (Figure 6A) and sodium oxamate (Figure 6B), confirming the coordinated regulation through immunoblotting. Consistent with protein-level observations, RT-qPCR analysis confirmed transcriptional regulation of AF9 in response to Kla perturbation (Figure 6C). Therefore, the changes in AF9 levels mirrored those of Kla and H3K9la, being upregulated by sodium L-lactate and downregulated by sodium oxamate. With prolonged treatment, AF9 levels gradually returned to baseline after Kla levels plateaued.

**Figure 6.**
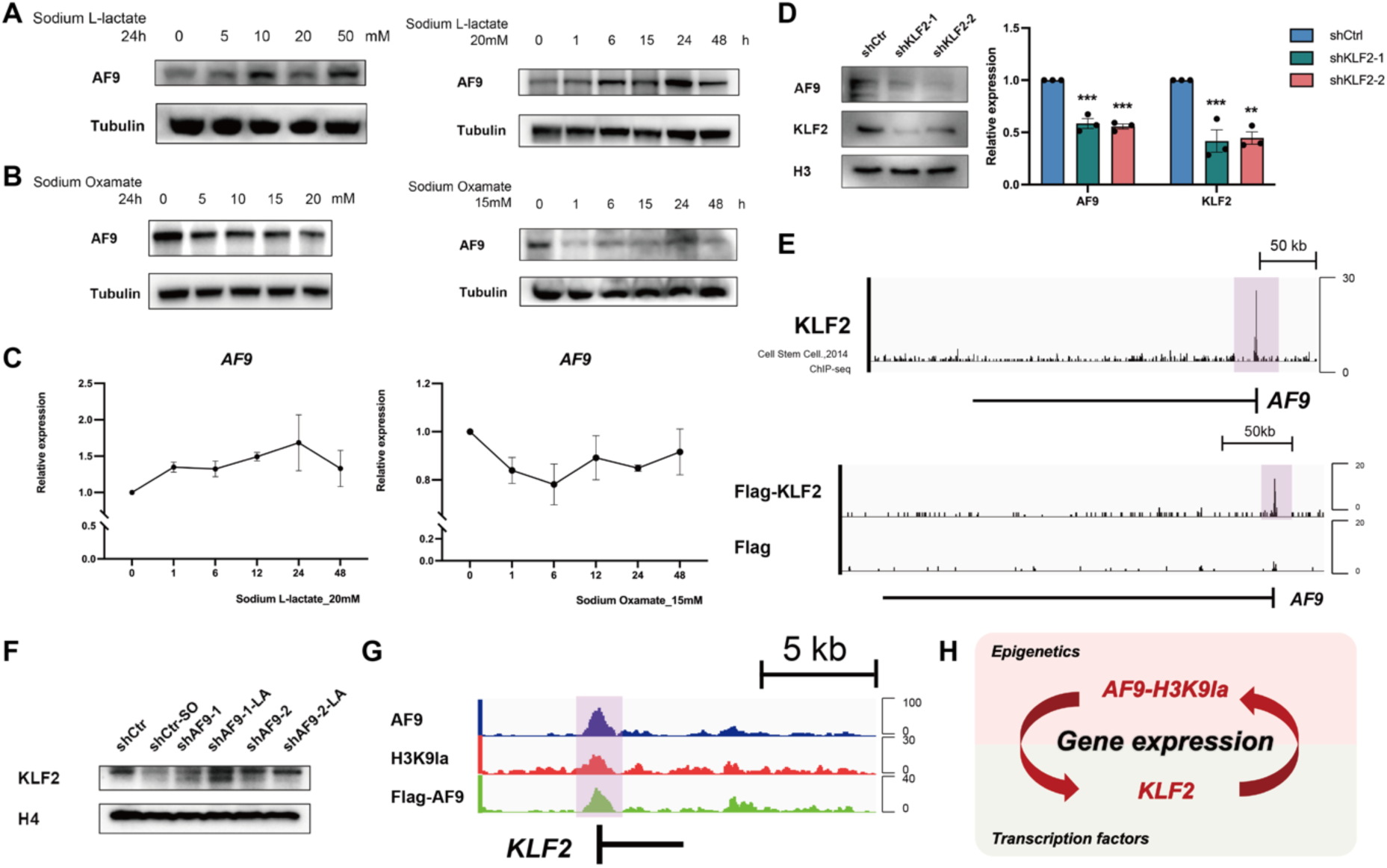
Positive feedback regulation of AF9 expression by H3K9la. (A, B) Western blot analysis of AF9 levels in response to the indicated concentrations of sodium L-lactate and sodium oxamate titration treatment in MCF7 cells. (C) RT-qPCR analysis of AF9 in MCF7 cells treated by sodium L-lactate and sodium oxamate. (D) The left is western blot analysis of AF9 and KLF2 in control (shControl) and KLF2 knockdown (shKLF2) MCF7 cells. Tubulin served as loading controls. The right is RT-qPCR analysis of AF9 and KLF2 in KLF2 knockdown (shKLF2) MCF7 cells. (E) Genome browser view of KLF2 peaks on AF9 gene promoter by reported data^23^ and Flag-KLF2 peaks occupied on AF9 promoter in MCF7 cells. (F) KLF2 is associated with the content of AF9 and Kla. (G) Genome browser view of AF9 (blue), H3K9la (red) and Flag-AF9 (green) CUT&Tag peaks on the indicated KLF2 gene in MCF7 cells. (H) “AF9-H3K9la/KLF2/AF9” positive feedback pathway.

We hypothesized that AF9 recognizes histone lactylation through a positive feedback mechanism. Initially, we proposed that transcription factors may serve as intermediaries in this process. To explore this, we predicted potential transcriptional regulators of AF9 and found that many belonged to the KLF family (Figure S7A). We screened for predicted KLF family members using spatial transcriptomics and identified KLF2 as being highly expressed in tumor cells (Figure S7B). Notably, KLF2 is frequently amplified in breast cancer, consistent with the amplification pattern observed for AF9 (Figures S7C and S7D). After analyzing the published ChIP-seq data of KLF2^32^, we detected that KLF2 was prominently positioned at the AF9 promoter (Figure 6E). To validate the co-localization in MCF7 cells, we repeated the CUT&Tag sequencing experiment (Figure 6E). Additionally, knockdown of KLF2 led to a reduction in AF9 expression (Figure 6D), suggesting that KLF2 acts as a transcription factor of AF9.

Intriguingly, we also found that AF9 and H3K9la co-localize at the promoter region of the KLF2 gene and appear to regulate its expression (Figure 6F and 6G). Moreover, the target genes of KLF2 and AF9 partially overlapped, suggesting that they may participate in a shared regulatory pathway (Figure S7E and S7F). These findings support the existence of a positive feedback loop in breast cancer, mediated by the interplay between epigenetic modification and transcriptional regulation involving AF9, H3K9la, and KLF2 (Figure 6H).

### AF9 is associated with the progression of breast cancer

Loss of AF9–H3K9la impairs the proliferation and invasive capacity of breast cancer cell lines. Analysis of TCGA data revealed that both AF9 and KLF2 are frequently amplified in breast cancer. These observations led us to hypothesize that elevated AF9 expression may be associated with increased tumor malignancy. Given these considerations, we performed an integrated analysis of spatial transcriptomics and RNA-seq data to characterize the AF9-associated spatial landscape of tumor cells within the tumor microenvironment (TME) of luminal breast cancer patient samples. AF9 exhibited strong co-localization with classified tumor cells, particularly within the tumor core (Figure 7A). Meanwhile, the scRNA-seq data enabled the visualization of multiple cell types within the sample (Figure 7B). Subsequently, we identified two tumor cell clusters distinguished by AF9 expression levels—high and low, respectively (Figure 7C). Consistent with previous findings, the AF9 high-expression cluster exhibited marked upregulation of the TGFβ signaling pathway, with additional enrichment of E2F targets and the EMT pathway (Figure 7D). We also observed a correlation between AF9 and its downstream target genes identified in the MCF7 cell line (Figure 7E).

**Figure 7.**
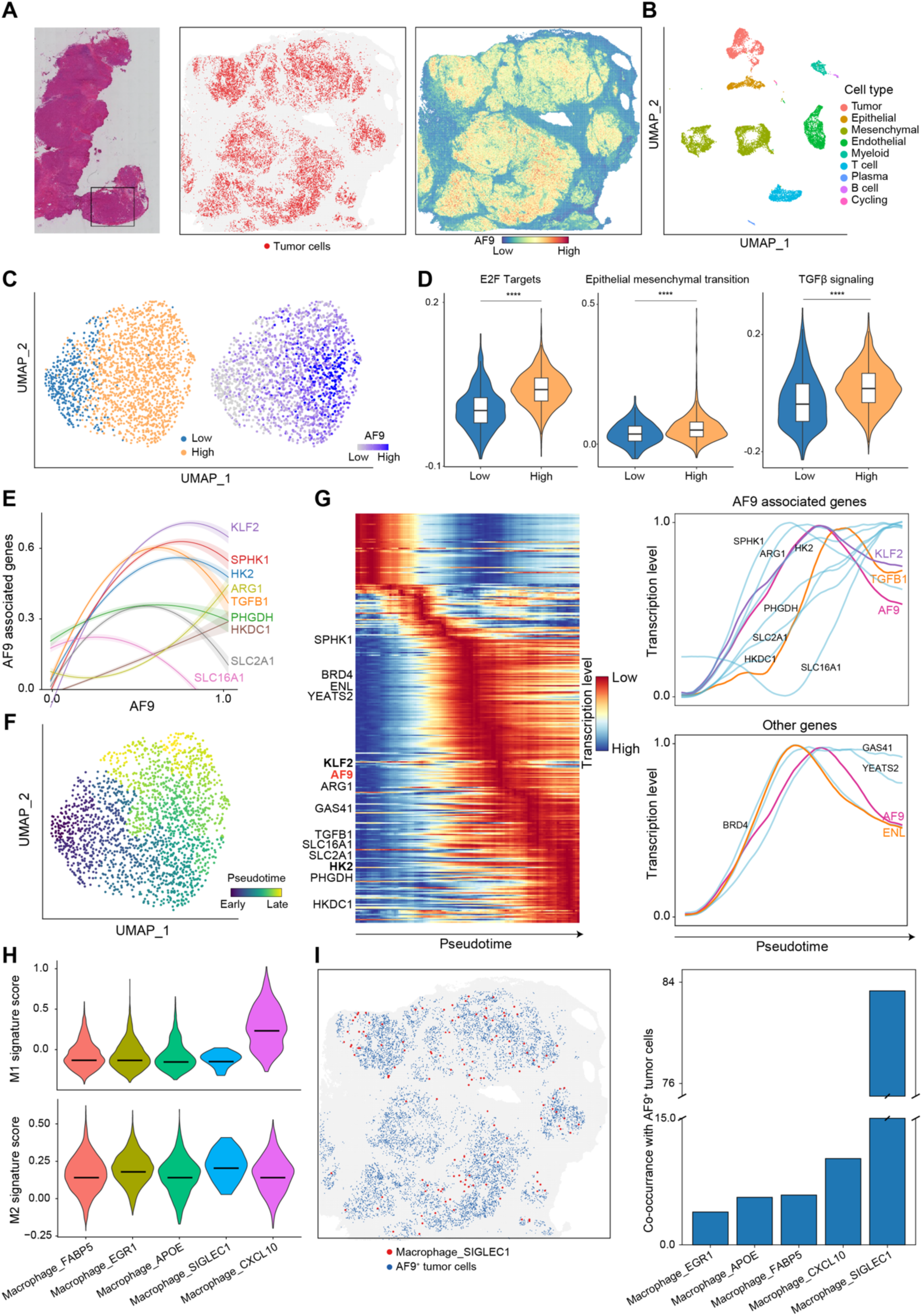
AF9 is associated with the progression of breast cancer. (A) Stereo-seq data of a breast cancer section, including the hematoxylin and eosin (H&E) staining (left panel), identified tumor bins (middle panel) and AF9 expression pattern (right panel). (B) Umap visualization showing the cell type from scRNA-seq data derived from a breast cancer sample. (C) Umap visualization illustrating two clusters of tumor cells (left panel) from scRNA-seq and corresponding expression of AF9 (right panel). (D) Violin plots showing the signature score of E2F targets, Epithelial mesenchymal and TGFβ signaling pathways in two clusters of tumor cells. Related to (C). (E) Line plot displaying the correlation of AF9 (x axis) and AF9 regulated genes (y axis) between the tumor cells in (C). (F) Umap visualization showing the pseudotime of tumor cells inferred by Slingshot. (G) Left panel: heatmap displaying expression profiles of differentially expressed genes along the differentiation pseudotime in (F). Right panel: line plots show the changes in expression profiles for AF9 regulated genes (top) and control genes (bottom) along the differentiation pseudotime, respectively. (H) Violin plots showing the signature score of M1 (top) and M2 (bottom) across different subtypes of macrophages. (I) Spatial plot in left panel displaying the distribution of Macrophage_SIGLEC1 and AF9^+^ tumor cells. Bar plot in right panel showing the spatial co-occurrance between AF9^+^ tumor cells with different subtypes of macrophages.

After reconstructing the pseudotime trajectory of tumor cells, we identified genes whose expression varied along the trajectory (Figure 7F). Interestingly, AF9 operated in a distinct temporal phase compared to other YEATS domain-containing reader proteins (Figure 7G), suggesting that its unique role in recognizing histone lactylation may contribute to its temporally distinct function. Notably, KLF2 expression slightly preceded that of AF9, implying a potential upstream regulatory role for KLF2 in modulating AF9 expression and subsequent tumor progression, as reflected in the line plot trends (Figure 7G).

Given that histone lactylation promotes the immunosuppressive activity of monocyte-derived macrophages^11,33^, and that AF9-mediated regulation of histone lactylation enhances glycolysis, we hypothesized that AF9 may interact with tumor-associated macrophages, particularly within the tumor microenvironment (TME). Supporting this, we observed that AF9-positive epithelial cells exhibited increased interactions with myeloid cells (Figure S6E). To further assess the spatial relevance of macrophage localization within tumors, we classified macrophages into inflammatory M1 and reparative M2 subtypes based on distinct marker expression (Figure 7H). Intriguingly, M2 macrophages were predominantly distributed around tumor cells with high AF9 expression (Figure 7I). These findings suggest that tumor cells with high AF9 expression may possess an enhanced ability to evade immune surveillance. In combination with the pro-tumorigenic signaling pathways enriched in AF9-high tumor cells, this immune-evasive capacity may contribute to tumor progression. Therefore, the AF9–H3K9la regulatory axis represents a potential therapeutic target in breast cancer.

## DISCUSSION

Lactate, a key glycolytic metabolite, has recently been recognized as a crucial link between metabolism and gene regulation via histone lactylation—an epigenetic modification implicated in macrophage polarization, cellular reprogramming, and tumor progression. Yet, how histone lactylation differs functionally from acetylation and contributes to gene-specific regulation remains poorly understood.

In this study, we identified the YEATS domain protein AF9 as a reader of histone lactylation. Through biochemical and genomic analyses, we demonstrated that AF9 binds H3K9la and promotes the expression of tumor-associated genes in breast cancer cells, enhancing invasiveness and modestly affecting proliferation. These genes are enriched in metabolic and signaling pathways, suggesting that AF9–H3K9la recognition drives oncogenic transcriptional programs.

Mechanistically, we uncovered a positive feedback loop in which AF9 activates the transcription factor KLF2, which in turn upregulates AF9 expression. Structural dissection using the AF9-F28R mutant further revealed that this regulation is largely dependent on lactylation rather than acetylation, particularly in the context of TGFβ signaling. TGFβ1 and histone lactylation have been implicated in promoting immunosuppressive TME features and enhancing HK2 of glycolysis, linking epigenetic and metabolic regulation^11,19,33–35^.

To validate this mechanism in the context of tumor microenvironment (TME), we integrated spatial transcriptomics and single-cell RNA-seq in luminal breast cancer samples. We observed that AF9-high tumor cells exhibit enhanced glycolytic activity, activate tumor-promoting pathways, and are frequently surrounded by immunosuppressive myeloid cells. Collectively, our findings reveal a multilayered regulatory circuit in which histone lactylation, mediated by AF9, orchestrates a ‘super’ positive feedback loop involving metabolism, transcription factors and signaling pathways. This network not only underlies tumor progression but also highlights the AF9–H3K9la axis as a promising therapeutic target in luminal breast cancer.

## METHOD DETAILS

### Materials

All histone peptides bearing different modifications (>98% purity) were synthesized at SciLight Biotechnology and summarized in Table S1. PTM antibodies including anti-H4 (PTM-1009), anti-pan Kla (PTM-1401RM), anti-H3K9la (PTM-1419RM) and anti-H3K18la (PTM-1406RM) were obtained from PTM Biolabs. Anti-AF9 (ab154492) and anti-Tubulin (ab231082) and anti-H3K9ac (ab32129) antibodies were purchased from Abcam; anti-FLAG (F1804) was purchased from Sigma; anti-PHGDH (bs-2970R) was purchased from Bioss; anti-TGFβ1 (D121324) was purchased from Sangon; and anti-KLF2 (TD13602S) was purchased from Abmart. Sodium L-lactate (Sigma, L7022) and sodium oxamate (ApexBio, C3893) were of commercial source. CUT&Tag was performed according to the manufacturer’s instructions (Novoprotein, N259). siRNAs pLKO shRNA constructs were purchased from Sigma. The shRNA sequences were summarized in Table S3.

### Protein expression and purification

The YEATS domains of AF9 (aa 1 to 138), ENL (aa 1 to 148) and GAS41 (aa 14 to 159) as well as the BRD domains of BRD3 (aa 306 to 416) and BRD9 (aa 131 to 250) were cloned into pET28b vector. The YEATS domain of YEATS2 (aa 201 to 332) as well as the DPF domains of DPF1 (aa 246 to 320), DPF2 (aa 270 to 391), DPF3 (aa 254 to 368), MOZ (aa 194 to 323) and PHF10 (aa 289 to 410) were cloned into pET28b vector with an N-terminal 10x His-SUMO tag. The DPF domain of MORF (aa 211 to 322) was cloned into pGEX-6P-1 vector.

All proteins were expressed in *E. coli* BL21 (DE3) and induced overnight by 0.2mM IPTG at 16°C in LB medium. Cells were harvested and disrupted by homogenizer in lysis buffer [YEATS domain proteins: 20mM Tris (pH 7.5), 500mM NaCl and 5% glycerol, 20mM Imidazole; DPF and BRD domain proteins: 20mM Tris (pH 7.5), 100mM NaCl and 5% glycerol, 20mM Imidazole]. Supernatants were run onto the HisTrap or GST column followed by extensive washing. His, His-SUMO and GST tagged proteins were digested by thrombin, ULP1 and PreScission protease. Samples were subjected to gel filtration on a Superdex 75 10/300 column using the AKTA Purifier system. Purified fractions were analyzed by Coomassie blue staining and pooled, concentrated, and stored at −80°C.

### Isothermal Titration Calorimetry

The titration was performed using the MicroCal PEAQ-ITC instrument (Malvern Instrument) at 15°C for YEATS domains as well as 25°C for DPF and BRD domains. Each ITC titration consisted of 17 successive injections. Usually, H3 peptides at 1.0-1.2mM were titrated into the recombinant proteins at 0.07-0.1mM. The resultant ITC curves were analyzed with Origin 7.0 (OriginLab) using the “One Set of Binding” fitting model. Protein concentrations were measured based on the UV absorption at 280nm.

### Crystallization, data collection, and structure determination

Crystallization was performed via the sitting drop vapor diffusion method at 16°C by mixing (0.2-1µl) protein with reservoir solution. The protein samples were prepared by mixing AF9-YEATS domain (aa 1 to 138, WT or F28R) with H3_1-10_K9la peptide in a molar ratio of 1:10 overnight at 4°C. The crystal of AF9_1-138_-H3K9la was grown in a reservoir solution containing 2.4M Sodium Malonate. And the reservoir solution of AF9-F28R_1-138_-H3K9la was 2.1M DL Malic acid pH7.0, 8% Ethylene glycol. The co-crystals were briefly soaked in cryoprotectant, comprised of reservoir solution supplemented with 15-25% ethylene glycol, and then flash-frozen in liquid nitrogen for data collection.

Diffraction data of crystals was collected at beamline BL02U at Shanghai Synchrotron Radiation Facility at 0.9792Å. Selected diffraction images were indexed, integrated, and merged using the HKL2000 software^36^. The structures were solved by molecular replacement using MOLREP^37^ from the CCP4 suite with AF9-H3K9ac (PDB ID: 4TMP) as the search model. Refinement and model building was performed with PHENIX^38^ and COOT^39^, respectively. The data collection and structure refinement statistics were summarized in Table S2. Structural figures were created using the PYMOL or ChimeraX programs.

### Cell culture, transfection and histone extraction

All cell lines were purchased from ATCC and tested for mycoplasma contamination (YEASEN, 40612ES25). MCF7, H1299 and HCT116 cell lines were maintained in RPMI 1640 medium (Gibco) supplemented with 10% fetal bovine serum (Gibco). HEK293T and HeLa cell lines was cultured in DMEM (Gibco) supplemented with 10% fetal bovine serum (Gibco). DNA and shRNA transient transfection were performed using Lipofectamine 3000 (Invitrogen) according to the manufacturer’s instructions. MCF7 was treated by sodium L-lactate and sodium oxamate and collected. Histone was extracted according to the epiQuik total histone extraction kit (Epigentek, OP-0006). Histone concentration was measured by the BCA method.

### Western blot analysis

Protein samples were separated on SDS polyacrylamide gel electrophoresis gels and then transferred to polyvinylidene fluoride (PVDF) membranes. The membranes were blocked in 5% nonfat milk diluted in TBST [20mM Tris (pH 8.0), 150mM NaCl, 0.1% Tween 20] for 1h at room temperature. The membranes were then incubated with primary antibodies at 4°C overnight. Then the membranes were washed with TBST and incubated with horseradish peroxidase conjugated anti-mouse or anti-rabbit antibodies used at a concentration of 1: 5,000 for 1h. Western blots were developed with an enhanced chemiluminescence detection system.

### RNA Extraction, Reverse Transcription, and Real-Time PCR Analysis

Total RNA was extracted using RNAsimple kit (Tiangen, DP419) and reverse-transcription used the All-in-one RT kit (Vazyme, R333). Quantitive real-time PCR (qPCR) analysis were performed using Power SYBR Green PCR Master Mix and the CFX96 Real-Time PCR Detection System (Bio-Rad). Gene expressions were calculated following normalization to ACTIN levels using the comparative CT (ΔΔCT) method. Statistic differences were calculated using a two-tailed unpaired Student’s test. The primer sequences for qPCR are listed in Table S4.

### Co-Immunoprecipitation

Cells were lysed in IP lysis buffer (Beyotime, P0013) supplemented with complete protease inhibitor tablet (Roche) followed by sonication. Anti-FLAG magnetic beads (Beyotime, P2115) were incubated with the lysates overnight at 4°C. The beads were then washed 3-5 times with IP lysis buffer, and the bound proteins were eluted in SDS-loading buffer and analyzed by Western blot.

### Targeted metabolomic analysis method

Targeted metabolomic experiment was analyzed by TSQ Quantiva (Thermo, CA). C18 based reverse phase chromatography was utilized with 10mM tributylamine, 15mM acetate in water and 100% methanol as mobile phase A and B respectively. This analysis focused on TCA cycle, glycolysis pathway, pentose phosphate pathway, amino acids and purine metabolism. In this experiment, we used a 25-minute gradient from 5% to 90% mobile B. Positive-negative ion switching mode was performed for data acquisition. The resolution for Q1 and Q3 are both 0.7FWHM. The source voltage was 3500v for positive and 2500v for negative ion mode. The source parameters are as follows: spray voltage: 3000v; capillary temperature: 320oC; heater temperature: 300oC; sheath gas flow rate: 35; auxiliary gas flow rate: 10. Metabolite identification was based on Tracefinder search with home-built database containing about 300 compounds.

### CUT&Tag data processing and analysis

The hyperactive Universal CUT&Tag Assay Kit for Illumina (Vazyme, TD903-1) was used for the test. Sequencing was conducted on a NovaSeq 6000 (Illumina) with paired-end 150-base reads. Alignment to the human (hg38) genome was performed using BWA (v0.7.15). Samtools (v1.3.1) was utilized for handling SAM and BAM files, and duplicates were removed using Picard (v1.107). BAM files were converted to bigWig format using bamCoverage from deepTools (v3.3.0) with RPKM normalization and a bin size of 10 bp. ENCODE blacklist regions were excluded prior to downstream analysis. Peak calling employed MACS2 (v2.1.2) with parameters: --nomodel --shift 100 --extsize 200.

Co-localization analysis of H3K9me3, AF9, and H3K9ac utilized the computeMatrix function of deepTools in reference-point mode, with parameters: -a 5000 -b 5000 --binSize 10 --sortUsingSamples 1 --skipZeros, and excluded blacklist regions. Genes with significant AF9 localization in promoters were identified by rank ordering AF9 binding density using featureCounts (v2.0.4).

### RNA-seq data analysis

Total RNAs were purified and high-throughoutput sequencing was performed by DNBSEQT7 (Beijing Genomics Institute, BGI). Sequencing reads were aligned to the human (hg38) genome using Hisat2 (v2.1.0). Reads with mapping quality <20 was filtered out using Samtools. Read counts were obtained using htseq-count from HTSeq (v2.0.3). Raw counts were processed using DESeq2 (v1.46.0) in R, defining differentially expressed genes as those with fold change >2 and adjusted p-value <0.05.

### ScRNA-seq data analysis

ScRNA-seq data analysis from either the in-house scRNA-seq dataset or Wu et al.’s scRNA-seq dataset obtained from GEO (https://www.ncbi.nlm.nih.gov/geo/, accession: GSE176078) were processed using the R package Seurat v4.0.5 in R v4.1.0 using the standard pipeline. Briefly, we performed log-normalization by using the NormalizeData function with a scale factor of 10,000 and identified highly variable features using the FindVariableFeatures function. For dimensional reduction, we used RunPCA with the first 50 principal components and performed graph-based clustering with a resolution of 0.5 using the FindClusters function. Batch correction was performed with the Harmony package using RunHarmony function. We then employed RunUMAP to execute uniform manifold approximation and projection (UMAP). Cell-cell communication was analyzed with the ‘CellPhoneDB’ package in python v3.7, the interaction was defined as the number of significant (p-value < 0.01) interacting gene pairs. Visualizations were generated and arranged in the ‘ggplot2’ package.

### Spatial Transcriptomics (ST) analysis

The ST libraries were prepared following the Stereo-seq protocol^40^. The libraries were sequenced on an MGI DNBSEQ-Tx sequencer and the RNAs were captured on each DNB spot with a resolution of 500 nm. Raw sequencing reads were filtered and demultiplexed using the SAW pipeline (https://github.com/BGIResearch/SAW). We generated a coordinate identity-containing expression profile matrix for each library. Gene expression levels in the ST dataset were projected onto the spatial image using the default functions with Seurat package.

### Enrichment analysis

The KEGG pathway enrichment analysis was performed using clusterProfiler (v4.14.4) package in R. The significant enriched pathways were visualized using circlize (v0.4.11) and dotplot function in clusterProfiler.

### Track visualization

Presentive track visualization was performed using the WashU Epigenome Browser, with bigWig files generated in the preceding steps. The visualization was optimized to display genomic tracks in a manner that allows for detailed inspection of chromatin modifications and transcription factor binding sites.

#### Data acquisition

The following raw data were utilized from publicly accessible databases: the alteration data of AF9 and KLF2 were retrieved from the cBioPortal for Cancer Genomics database^41^. All datasets generated in this study have been deposited in the NCBI Gene Expression Omnibus under accession number <u>GSE291224 and GSE291225</u>.

#### TF prediction

Promoter regions were defined as extending 2000 bp upstream and 500 bp downstream of the transcription start site (TSS). Motif searching was performed using the FIMO tool (v5.5.7) from the MEME Suite. Motifs were identified based on a stringent *p*-value threshold of 0.0001 to ensure robust filtering and minimize false positives. The identified motifs were subsequently used for transcription factor (TF) binding site prediction, enabling the elucidation of potential regulatory elements within the defined promoter regions.

## RESOURCE AVAILABILITY

### Lead contact

Further information and requests for resources and reagents should be directed to and will be fulfilled by the lead contact, Haitao Li.

### Materials availability

All materials generated in this study are available from the lead contact with a completed Materials Transfer Agreement.

### Data and code availability

The structure data have been deposited at PDB.

All CUT&Tag and RNA-seq data have been deposited at GSE291224 and GSE291225.

All spatial transcriptomic sequencing data generated by Stereo-seq have been deposited at CNGB Sequence Archive (CNSA) of China National GeneBank DataBase (CNGBdb).

Any additional information required to reanalyze the data reported in this paper is available from the lead contact upon request.

## SUPPLEMENTAL INFORMATION

**Supplementary Figure S1.**
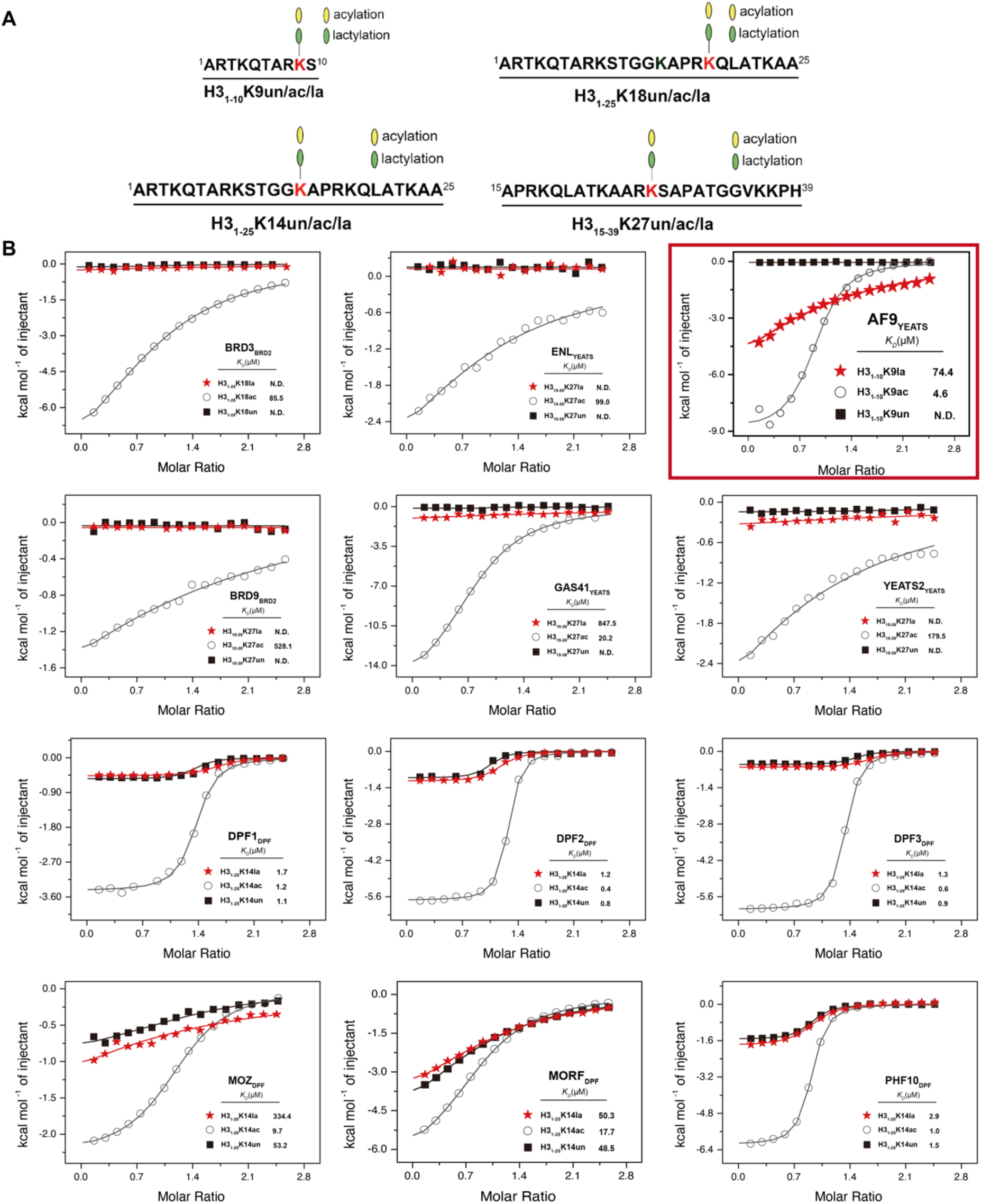
Recognition of histone lactylation by acylation reader proteins. (A) Peptide sequence synthesized in vitro. (B) ITC screen titration curve of histone acylation readers for non-modified (black), histone acetylation (brown), and histone lactylation (red) peptides.

**Supplementary Figure S2.**
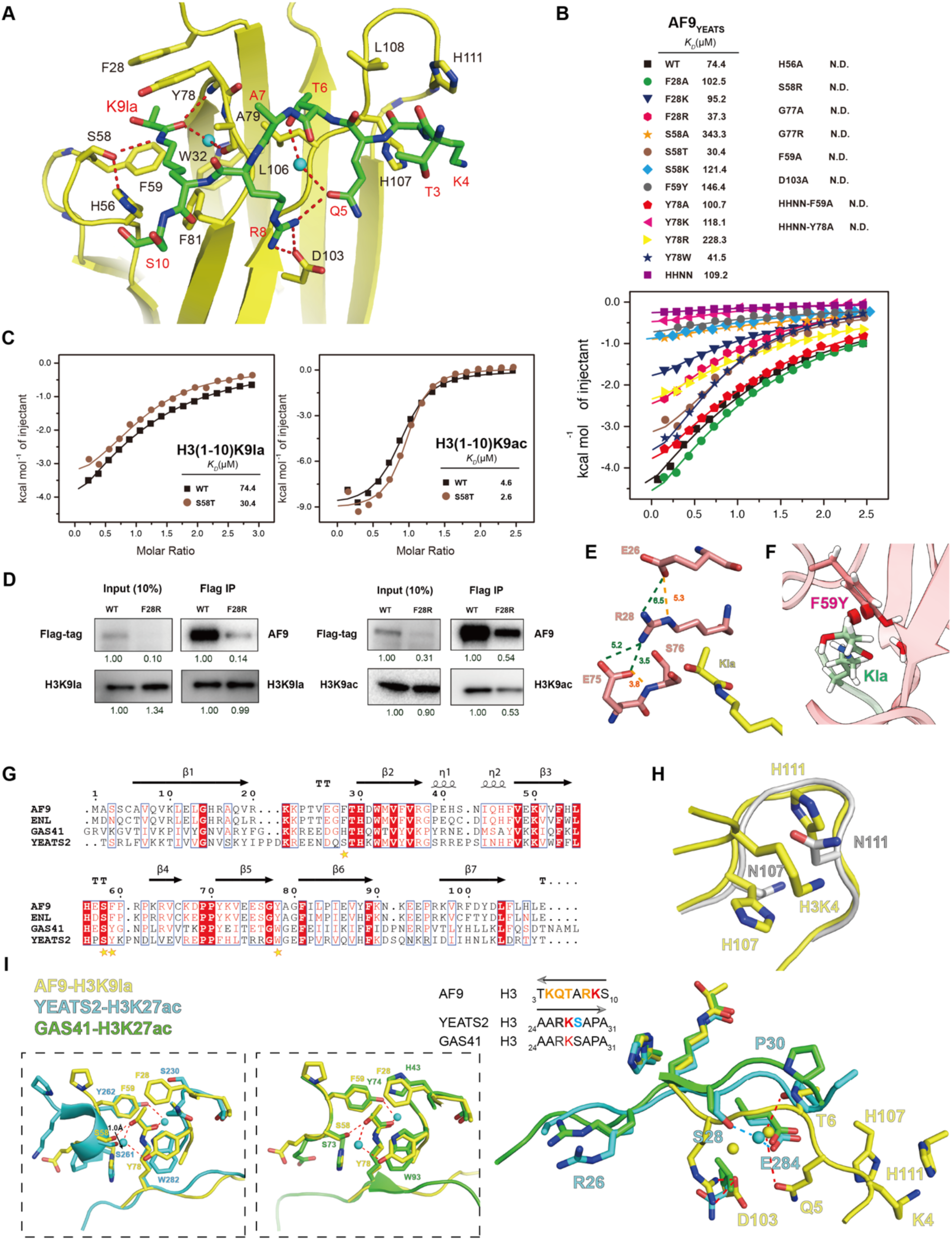
The AF9-F28R YEATS domain is capable of recognizing H3K9la through a mechanism that enhances its ability. (A) Stereo view of hydrogen bonding network involving H3 side chain (green sticks) and residues in AF9 YEATS (yellow sticks). Red dotted lines, hydrogen bonds; Cyan balls, water molecules. (B) Mutagenesis and ITC using mutant and WT AF9 YEATS with H3K9la peptide. (C) ITC assays of H3K9la (left) and H3K9ac (right) binding by WT or S56T mutant of AF9 YEATS. (D) Coimmunoprecipitation analysis of the interaction of Flag-AF9 WT or F28R mutant and H3K9la/H3K9ac. (E) The “polar gate” formed by electrostatic interaction and hydrogen bond interaction. Distances measured between atoms or centroids are color-coded in green or orange. (F) Close contact analyses of F59Y mutant structure bound to H3K9la. Red disk, van der Waals overlap. (G) Sequence alignment of 4 human YEATS domains. (H) Structural alignments of the reader pocket loop around AF9 YEATS and ENL YEATS (PDB ID: 5J9S) domains, which target Lys4 of H3 peptide. The key residues of AF9YEATS (yellow) and ENL YEATS (grey) domains are showed as sticks. (I) Structural alignments of the reader pocket loop around AF9 YEATS domian, YEATS2 YEATS (PDB ID: 5XNV) domain and GAS41 YEATS (PDB ID: 5XTZ), the key residues of AF9 YEATS (yellow), YEATS2 YEATS (blue) and GAS41 YEATS (green) domains are showed as sticks.

**Supplementary Figure S3.**
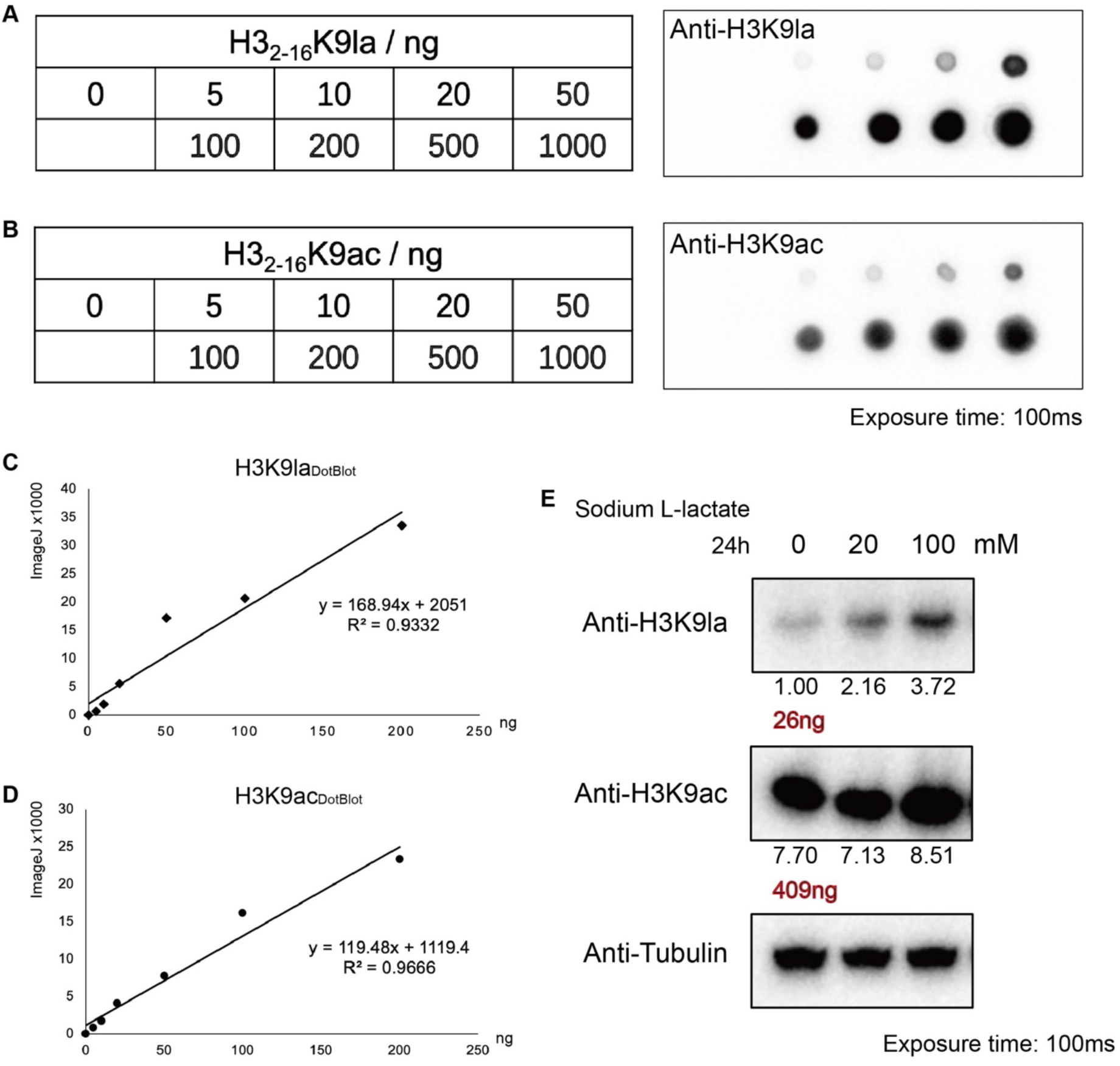
Quantification of H3K9la compared with H3K9ac. (A-D) Dot blot analysis of H3K9la and H3K9ac peptides and correspon€g fitting curves. (E) Western blot analysis of H3K9la and H3K9ac concentrations in MCF7 cells. Quantification of band intensity is performed using ImageJ.

**Supplementary Figure S4.**
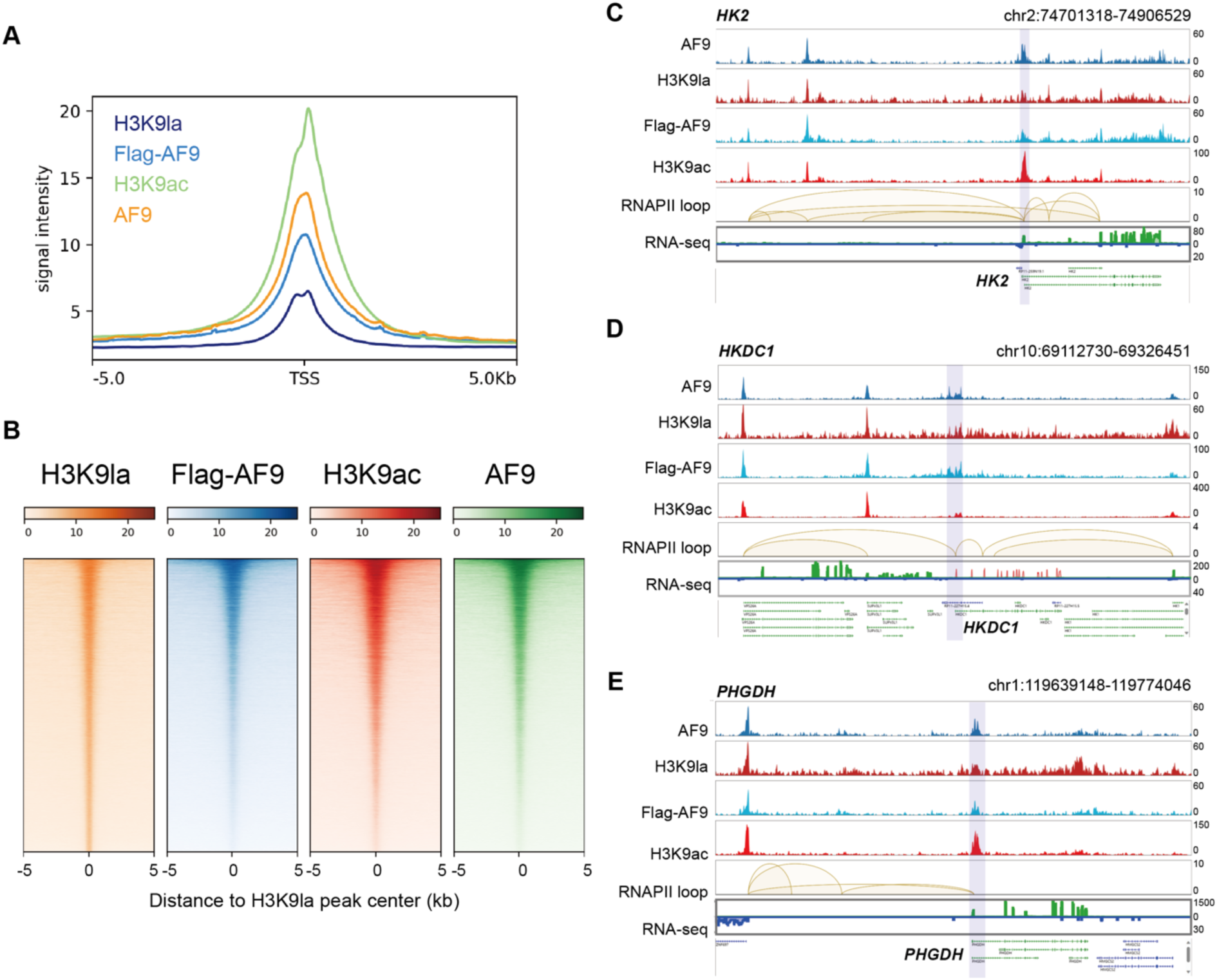
Co-localization of AF9, H3K9la and H3K9ac. (A-B) Metaplots and heatmaps depicting the co-localization of AF9, H3K9la and H3K9ac in gene promoters (A) and H3K9la peaks (B). (C-E) Representative screenshots showing co-localization of AF9, H3K9la and H3K9ac. The RNAPII ChIA-PET loop (GSM970209) and stranded-specific RNA-seq data (ENCFF301XEH: plus strand, ENCFF162JXM: minus strand) indicating active gene transcription-related chromosomal interaction and transcriptional state, support the reliability of the co-localization over HK2, HKDC1 and PHGDH gene promoters.

**Supplementary Figure S5.**
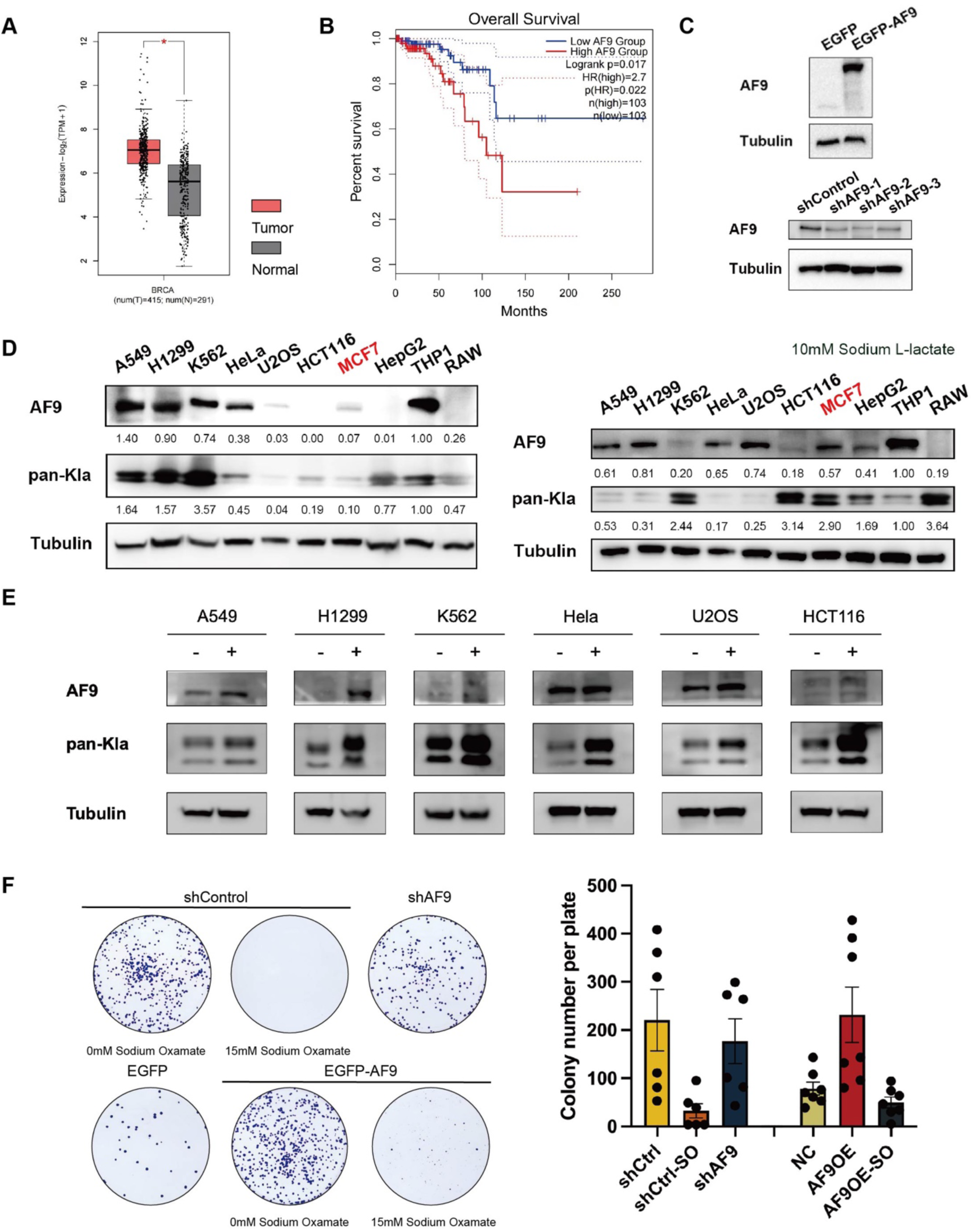
AF9 is associated with the development of luminal breast cancer. (A) AF9 is amplified in luminal A breast cancer. Data was acquired from TCGA. (B) AF9 is associated with poor prognosis in luminal A breast cancer patients. Data was acquired from TCGA. (C) Western blot analysis of AF9 knockdown (left) and overexpression (right) in MCF7 stable cell lines. Tubulin is used as loading controls. (D and E) Western blot analysis of AF9 and Kla concentration in cancer and macrophage cell lines with or without sodium L-lactate treatment. Tubulin used as the loading control. (F) AF9 is required for MCF7 cell proliferation (left) and analyzed by ImageJ (right). Colony formation assays of cells were stained with crystal violet 10 d after seeding.

**Supplementary Figure S6.**
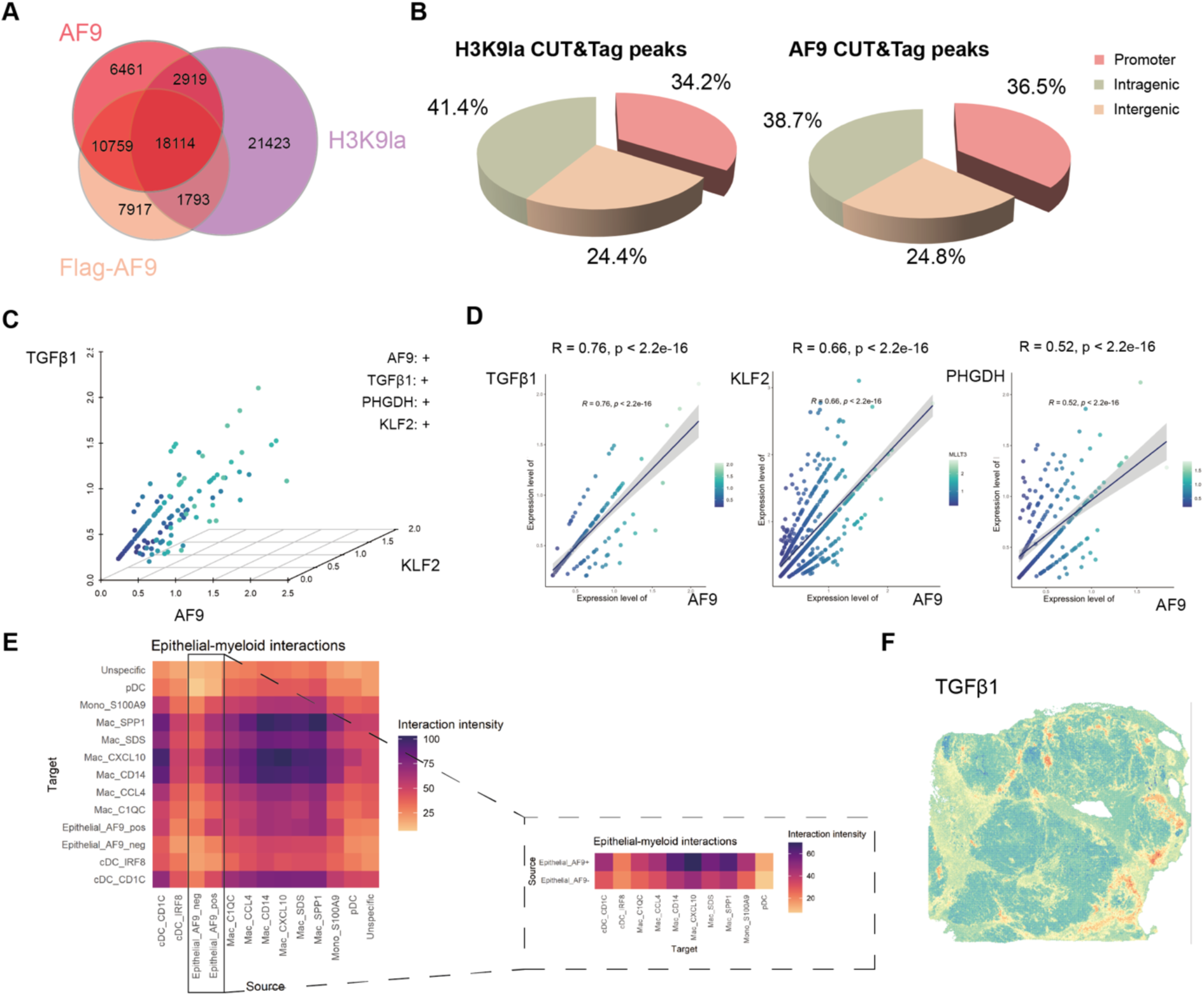
AF9 recognizes H3K9la promoting TGFβ signaling pathway and facilitating the establishment of intracellular communication. (A) Venn diagram showing the overlap of AF9, Flag-AF9 and H3K9la occupied peaks. (B) Genomic distribution of H3K9la (left) and AF9 (right) CUT&Tag peaks in MCF7 cells. The peaks are enriched in the promoter regions (Transcription Start Site [TSS] −/+5K). (C-D) Gene correlation analysis of AF9 as well as TGFβ1, PHGDH and KLF2. (E) Epithelial-myeloid cells interaction analysis. (F) Stereo-seq data of a breast cancer section, including the TGFβ1 expression pattern.

**Supplementary Figure S7.**
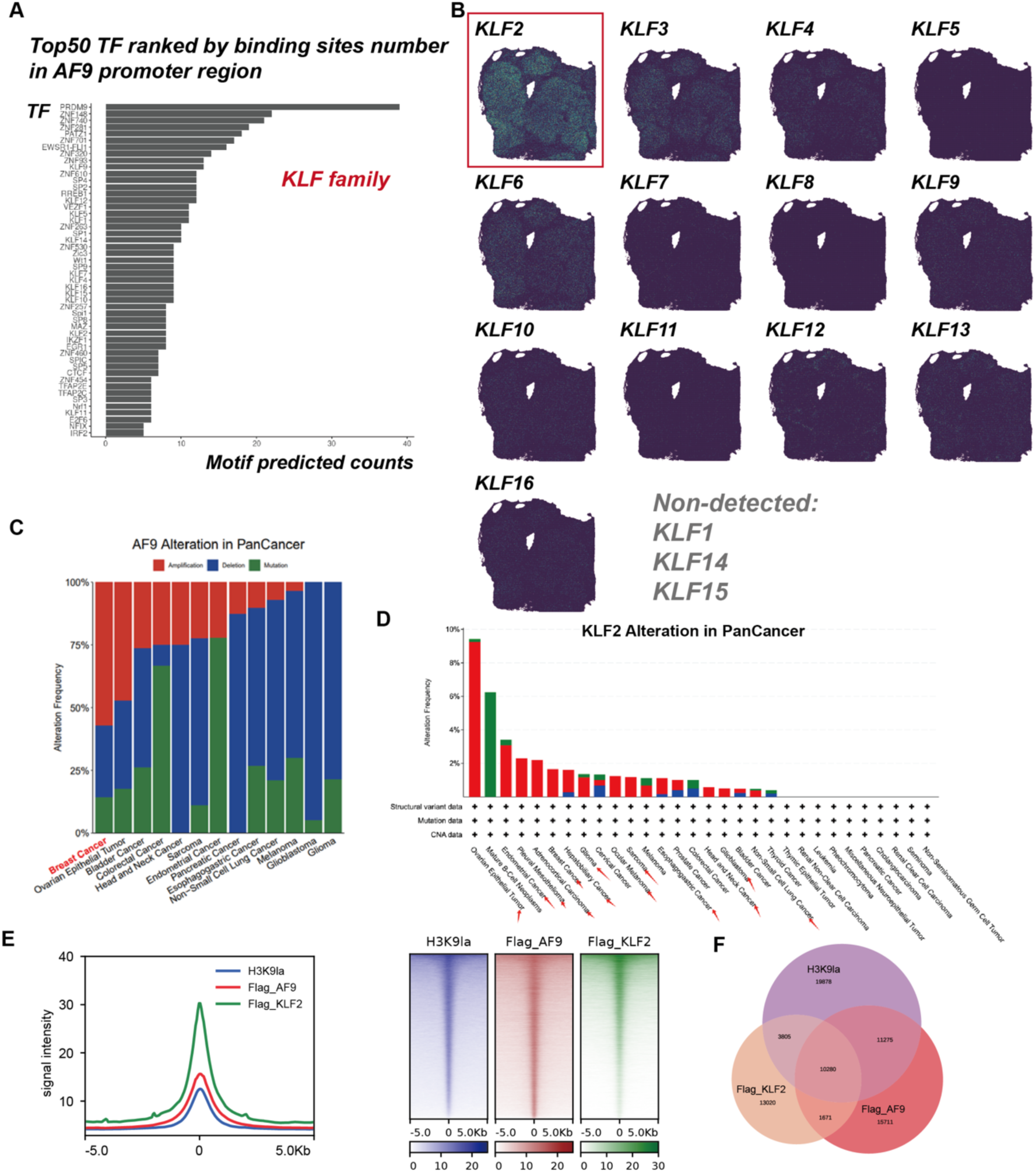
KLF2 is abundantly expressed in luminal breast cancer. (A) The anticipated transcription factors of AF9 gene. The significantly enriched TFs were ranked by their binding site number. (B) Predicted KLF family members in spatial transcriptomics data of a luminal breast cancer sample. (C) Histogram showing the proportion of different alteration frequency of AF9 in human cancers. Data was acquired from cBioPortal. (D) Histogram showing the alteration frequency of KLF2 in human cancers. Data was acquired from cBioPortal. (E) Average genome-wild occupancies (left) and heat maps of the normalized density (right) of Flag-AF9 (red), Flag-KLF2 (green) and H3K9la (violet) in MCF7 cells.

**Supplementary Table S1.** Layout of the histone peptides.

| Peptides | Sequence |
| --- | --- |
| H3<sub>1-10</sub>K9la | ARTKQTAR**Kla**S |
| H3<sub>1-10</sub>K9ac | ARTKQTAR**Kac**S |
| H3<sub>1-10</sub> | ARTKQTARKS |
| H3<sub>2-16</sub>K9la | RTKQTAR**Kla**STGGKAP |
| H3<sub>2-16</sub>K9ac | RTKQTAR**Kac**STGGKAP |
| H3<sub>1-25</sub>K14la | ARTKQTARKSTGG**Kla**APRKQLATKAA |
| H3<sub>1-25</sub>K14ac | ARTKQTARKSTGG**Kac**APRKQLATKAA |
| H3<sub>1-25</sub>K18la | ARTKQTARKSTGGKAPR**Kla**QLATKAA |
| H3<sub>1-25</sub>K18ac | ARTKQTARKSTGGKAPR**Kac**QLATKAA |
| H3<sub>1-25</sub> | ARTKQTARKSTGGKAPRKQLATKAA |
| H3<sub>15-19</sub>K18la | APR**Kla**Q |
| H3<sub>15-39</sub>K27la | APRKQLATKAAR**Kla**SAPATGGVKKPH |
| H3<sub>15-39</sub>K27ac | APRKQLATKAAR**Kac**SAPATGGVKKPH |
| H3<sub>15-39</sub> | APRKQLATKAARKSAPATGGVKKPH |

**Supplemental Table S2.**
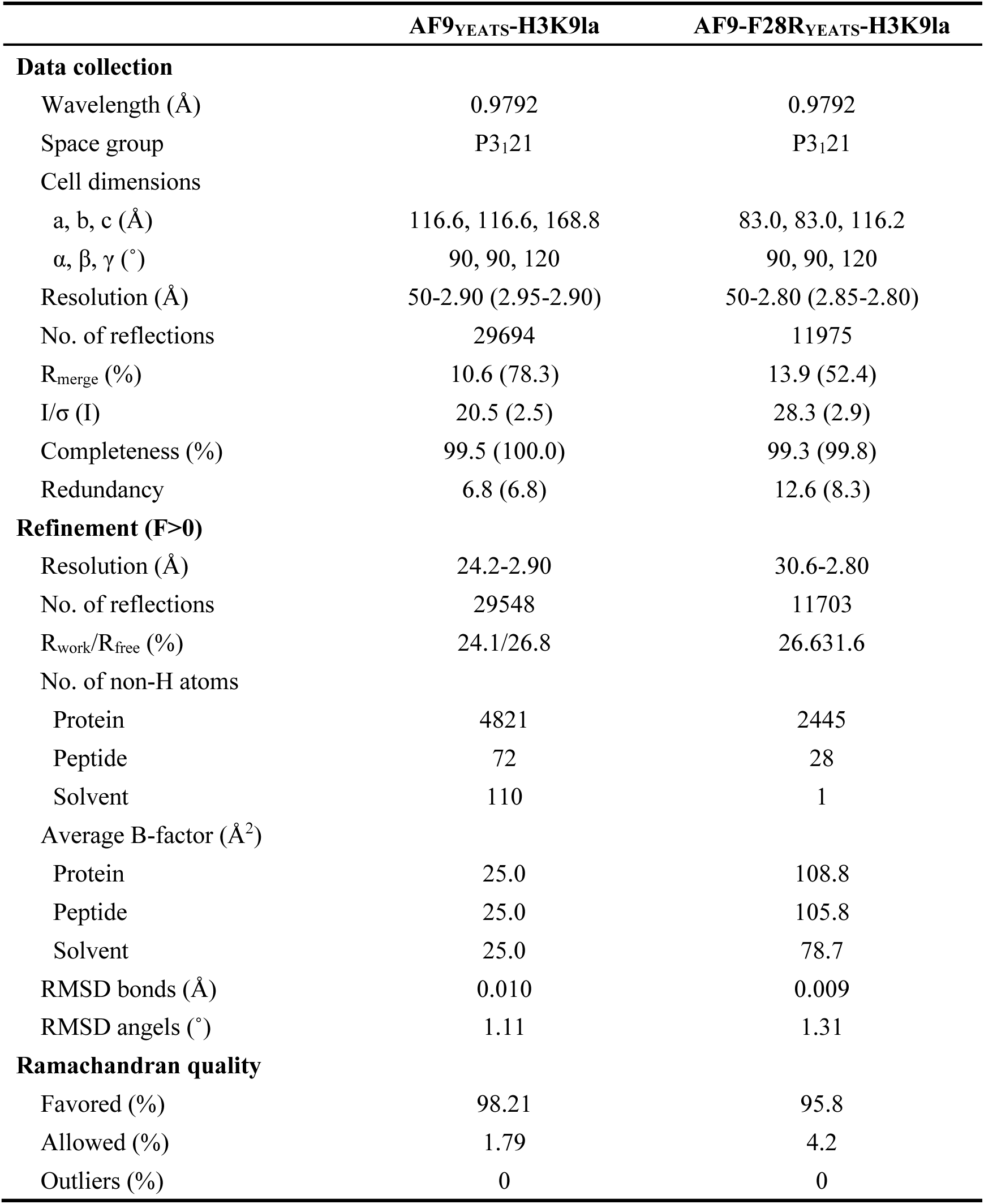
Data collection, processing and refinement statistics.

|  | **AF9<sup>YEATS</sup>-H3K9la** | **AF9-F28R<sup>YEATS</sup>-H3K9la** |
| --- | --- | --- |
| **Data collection** |  |  |
| Wavelength (Å) | 0.9792 | 0.9792 |
| Space group | P3<sub>1</sub>21 | P3<sub>1</sub>21 |
| Cell dimensions |  |  |
| a, b, c (Å) | 116.6, 116.6, 168.8 | 83.0, 83.0, 116.2 |
| $\alpha$, $\beta$, $\gamma$ (°) | 90, 90, 120 | 90, 90, 120 |
| Resolution (Å) | 50-2.90 (2.95-2.90) | 50-2.80 (2.85-2.80) |
| No. of reflections | 29694 | 11975 |
| R<sub>merge</sub> (%) | 10.6 (78.3) | 13.9 (52.4) |
| I/$\sigma$ (I) | 20.5 (2.5) | 28.3 (2.9) |
| Completeness (%) | 99.5 (100.0) | 99.3 (99.8) |
| Redundancy | 6.8 (6.8) | 12.6 (8.3) |
| **Refinement (F>0)** |  |  |
| Resolution (Å) | 24.2-2.90 | 30.6-2.80 |
| No. of reflections | 29548 | 11703 |
| R<sub>work</sub>/R<sub>free</sub> (%) | 24.1/26.8 | 26.6/31.6 |
| No. of non-H atoms |  |  |
| Protein | 4821 | 2445 |
| Peptide | 72 | 28 |
| Solvent | 110 | 1 |
| Average B-factor (Å<sup>2</sup>) |  |  |
| Protein | 25.0 | 108.8 |
| Peptide | 25.0 | 105.8 |
| Solvent | 25.0 | 78.7 |
| RMSD bonds (Å) | 0.010 | 0.009 |
| RMSD angels (°) | 1.11 | 1.31 |
| **Ramachandran quality** |  |  |
| Favored (%) | 98.21 | 95.8 |
| Allowed (%) | 1.79 | 4.2 |
| Outliers (%) | 0 | 0 |

**Supplemental Table S3.**
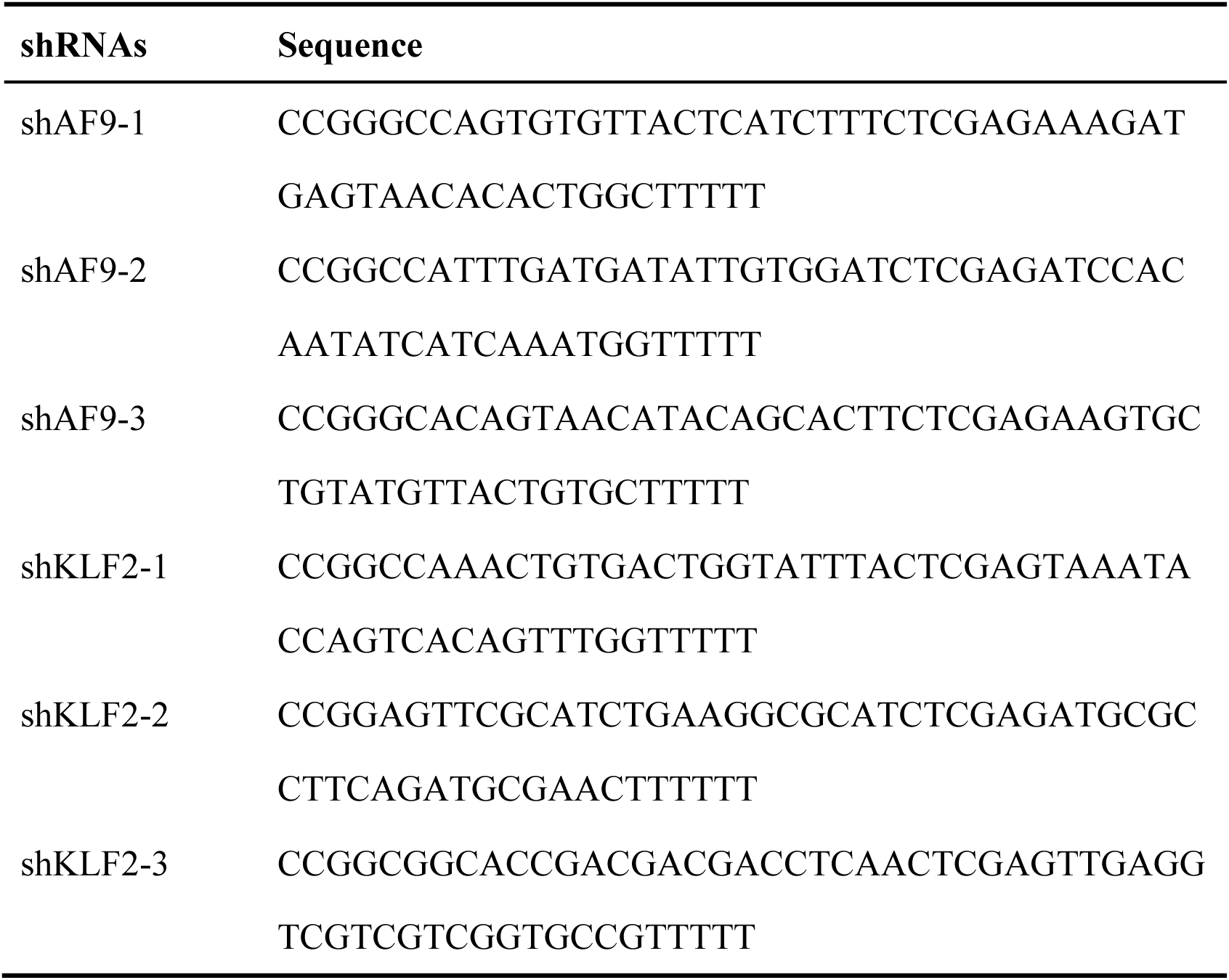
List of shRNAs.

**Supplemental Table S4.**
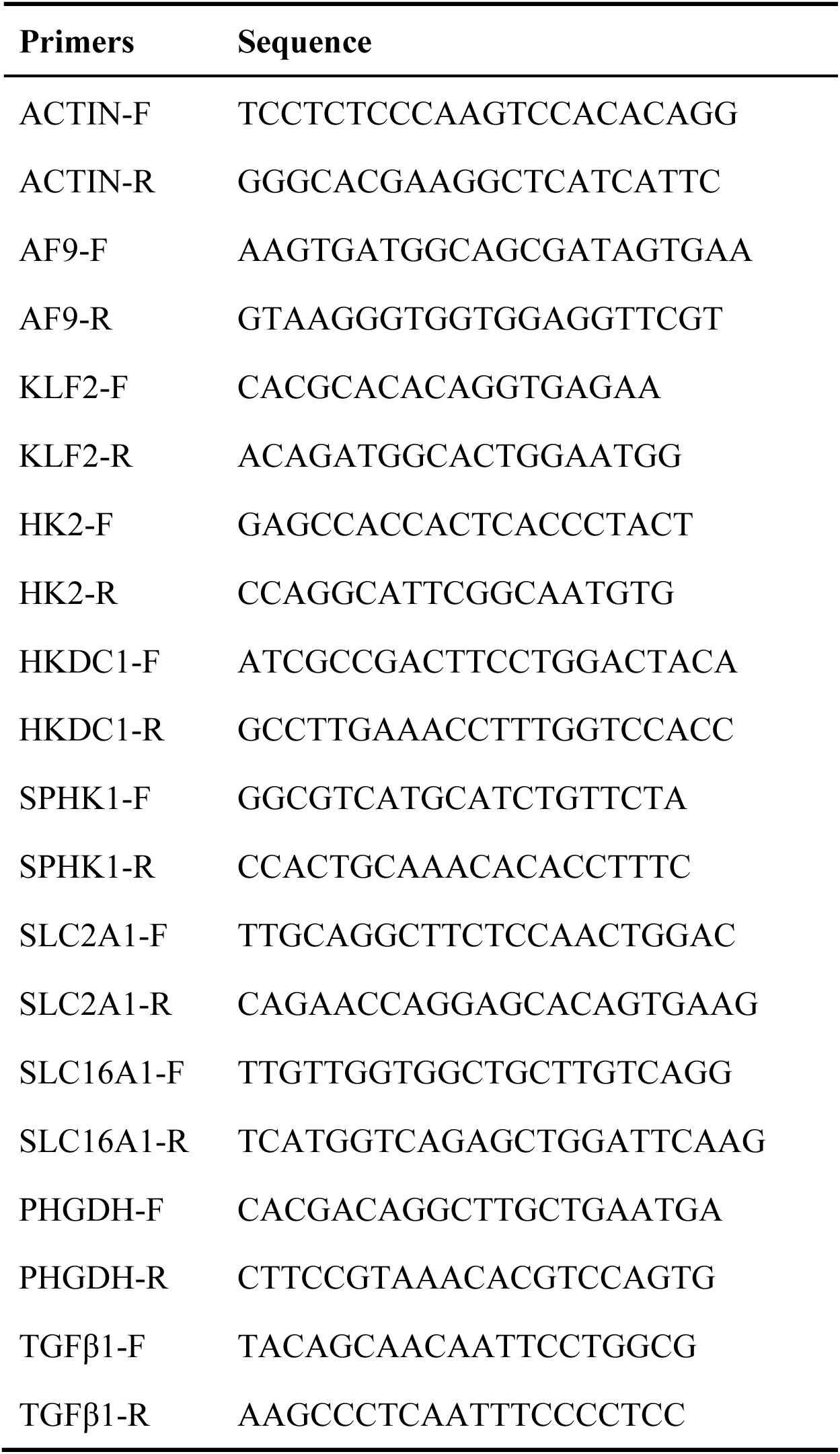
List of qPCR primers.

## ACKNOWLEDGMENTS

We thank other members in the Li laboratory for scientific inputs. We thank the staff members at beamline BL02U1 of the Shanghai Synchrotron Radiation Facility and S. Fan at Tsinghua Center for Structural Biology for assistance in data collection. We thank the staff of the Facility Center of Metabolomics and Lipidomics of China National Center for Protein Sciences at Tsinghua University. We thank Jiaxiang Liu from the Cancer Hospital, Chinese Academy of Medical Sciences & Peking Union Medical College for their assistance in tumor sample extraction. This research was supported by funding from the National Key R&D Program of China (2020YFA0803300 to H.L. and Z.T), National Natural Science Foundation of China (92153302 to H.L., 82373060 to X.W., 82203688 to C.Y.), Beijing Natural Science Foundation (JQ23032 to X.W.), CAMS Innovation Fund for Medical Sciences (CIFMS) (2021-I2M-1-014, 2023-I2M-C&T-A-009 to X.W.), Noncommunicable Chronic Diseases-National Science and Technology Major Project (2023ZD0502200 to X.W).

## AUTHOR CONTRIBUTIONS

H.L. conceived the study. H.L., Z.T., and X.W. supervised the studies. H.M. performed biochemical, structural and functional experiments. Z. Tang, M. Yuan and Y.Y. performed bioinformatic analyses. X. W. and C.Y. performed spatial transcriptomic studies of patient samples. H. L. and H. M wrote the manuscript with inputs from other authors.

## ACCESSION NUMBERS

The atomic coordinates and structure factors have been deposited in the Protein Data Bank under accession numbers 8Z73 (AF9 YEATS domain and H3K9la complex) and 9IM4 (AF9 YEATS domain F28R mutant and H3K9la complex). The CUT&Tag data (accession number GSE291224) and the RNA-seq data (accession number GSE291225) have been meticulously curated and deposited in the Gene Expression Omnibus (GEO) database. All spatial transcriptomic sequencing data generated by Stereo-seq have been deposited in the CNGB Sequence Archive (CNSA) of the China National GeneBank DataBase (CNGBdb) under accession number CNP0004319 and in the Open Archive for Miscellaneous Data (OMIX) under accession number OMIX004167.

